# Microgels enable iPSCs to assemble, expand, and differentiate into organoids – from sizable to high throughput

**DOI:** 10.1101/2025.10.20.683388

**Authors:** Laura Klasen, Matthias Mork, Ramin Nasehi, Vasudha Turuvekere Krishnamurthy, Kira Zeevaert, Aaron Babendreyer, Jacopo Di Russo, Wolfgang Wagner, Laura De Laporte

**Affiliations:** Institute for Technical and Macromolecular Chemistry, RWTH Aachen University, Worringerweg 1–2, 52074 Aachen, Germany; Department of Advanced Materials for Biomedicine (AMB), CBMS—Center for Biohybrid Medical Systems, AME—Institute of Applied Medical Engineering, RWTH Aachen University, Forckenbeckstraße 55, 52074 Aachen, Germany; DWI—Leibniz Institute for Interactive Materials e. V., Forckenbeckstrasse 50, 52074 Aachen, Germany; Helmholtz-Institute for Biomedical Engineering, RWTH Aachen University, Pauwelsstraße 20, 52074, Aachen, Germany; Institute for Stem Cell Biology, RWTH Aachen University Medical School, Pauwelsstraße 20, 52074, Aachen, Germany; Institute of Molecular and Cellular Anatomy, RWTH Aachen University, Wendlingweg 2, 52074 Aachen, Germany; Institute of Molecular Pharmacology, University Hospital RWTH Aachen University, Wendlingweg 2, 52074 Aachen, Germany

## Abstract

Organoid research holds tremendous potential for personalized medicine and drug development. However, current limitations include reproducibility issues largely due to the use of biologically derived materials, which are prone to batch-to-batch variations. Here, we report a new technology for human induced pluripotent stem cell (iPSC)-based organoid production with iPSC expansion and differentiation in the same construct in a reproducible and scalable manner, compatible with high-throughput automation. Chemically defined poly(ethylene glycol) (PEG)-based microgels are produced via parallelized step-emulsification microfluidics, enabling scalable production. This approach leverages the self-organization of iPSCs with microgels to build three-dimensional constructs, driven by robust cell-material interactions achieved through vitronectin-coated PEG microgels. This technology allows the iPSCs to expand and retain their pluripotency, after which they can be differentiated into the three germ layers, providing a suitable platform for organoid differentiation. This was further extended by differentiation into cardiac organoids and retinal photoreceptors to demonstrate two exemplary tissues.

## Main

Organoids are three-dimensional *in vitro* organ model systems of increasing interest in biomedical research for disease modeling and next-generation drug screening and development[2]. Due to their derivation from either pluripotent stem cells or organ-specific progenitor cells, organoids are developed for an increasing number of organs, including the liver[3–5], heart[6–8], brain[9–11], and retina[12, 13]. Induced pluripotent stem cells (iPSCs) are somatic cells reprogrammed to a pluripotent state through the forced expression of specific transcription factors, enabling self-renewal and differentiation into all germ layers. They provide a more ethically sustainable and accessible alternative to embryonic stem cells (ESCs)[14]. They have great potential in personalized medicine and organoid research and are receiving increasing attention[15].

In addition to advances in cell types, organoid research has also progressed by introducing various biomaterials that form scaffolds to enhance the cellular organization, stability, and uniformity[16], and the functionality of the resulting organoids[17–19]. Hydrogels that mimic the extracellular matrix (ECM), e.g., Matrigel[20], collagen[21, 22], fibrin[23, 24], hyaluronic acid (HA)[25], or decellularized tissue[26–28] are the main biomaterials used in organoid research. Since these biologically derived materials persistently demonstrate high batch-to-batch variability and could have tumorigenic potential[20, 29], the necessity arises for chemically well-defined biomaterial systems for organoid research[29–31], with one of the most frequently utilized synthetic hydrogels being poly(ethylene glycol) (PEG)[32, 33].

One main challenge in 3D tissue models is to grow dense functional tissue due to the reduced proliferation of many mature cells and insufficient diffusion. On the other hand, iPSCs have a high expansion potential but start to differentiate in an uncontrolled manner inside 3D hydrogels due to the many signals provided by the materials that highly influence cellular processes[34]. Therefore, a 3D hydrogel system that allows for iPSC expansion into dense cellular constructs and sequential controlled differentiation in the same construct can bridge this gap. Previous work has demonstrated that maintaining the self-renewal hallmark of the stem cell phenotype and, thereby, the proliferation capacity, in 3D hydrogels can be promoted by enhancing cell-cell interaction by the use of very dynamic[35] or fast degradable[36] hydrogels or the addition of single-cell survival-enhancing supplements [37].

In recent years, microporous annealed particle (MAP) scaffolds have emerged to enhance cell growth and infiltration by providing more empty space. MAP scaffolds are 3D constructs assembled by microgels and exhibit pore interconnectivity with micron-scale pores compared to conventional nanoporous hydrogels that must be degraded to provide sufficient space for the cells. Therefore, MAP scaffolds offer an alternative for bulk hydrogels that rely on an intricate balance between degradability and mechanical support to enable tissue formation [38–42]. To produce monodisperse spherical microdroplets and -gels at high production rates, parallelized microfluidic step-emulsification has emerged as a suitable method compared to single-channel microfluidic systems[43, 44].

Here, we demonstrate that iPSC-induced assembly of microgels resulted in millimeter-scale or high-throughput scaffolds that support iPSC growth while maintaining their stemness, allowing for sequential controlled differentiation at defined time points. Scalability of the microgel building block production is crucial for meeting the demands of growing larger organoids and high-throughput applications. We present a new method to grow human tissues *in vitro* in a reproducible and scalable manner by combining iPSC-based organoid technologies with microgel-based biomaterials in a chemically defined system. We hypothesized that this would be possible due to the high level of cell-cell interactions that this method allows for, in addition to facilitating diffusion through the microgel networks. Our scalable strategy is based on self-organized scaffolds built by iPSCs using functionalized PEG-based spherical microgels, yielding 3D constructs that allow for self-controlled expansion and guided differentiation. Microdroplets composed of PEG-diacrylate (PEG-DA) and glycidyl methacrylate (GMA) were fabricated by parallelized step-emulsification in a microfluidic device. Conversion into microgels is achieved by photoinitiated free-radical polymerization in-flow. The microgels were post-modified with vitronectin by amine-epoxy coupling to the epoxide moieties of the GMA incorporated into the polymer matrix. To demonstrate the large versatility and relevance of this system, sequential differentiation into the three germ layers is shown, and iPSC-derived cardiac organoids and retinal cells were generated. Additionally, we established the scalability and compatibility of our approach with an automated liquid handling system, highlighting its suitability for high-throughput production necessary for compound screening during drug development and disease modeling.

### Large-scale microfluidic production and biofunctionalization of spherical PEG-based microgels

To generate sufficient quantities of functionalized spherical microgels for iPSC-induced 3D assembly, we developed a parallelized step-emulsification microfluidic device (Fig. 1a). The dispersed phase, carrying the microgel precursors, is transferred through individual microchannels, and droplet formation is achieved by expansion into the deep oil reservoir. Through parallelization of these microchannels, we generated microdroplets with an average diameter of ∼ 80 µm, achieving a production rate in the range of 10^7^ droplets per hour (Supplementary Fig. 1 and Supplementary Video 1). After generation, the droplets were transferred off-chip through an outlet tube and photo- crosslinked in the outlet tubing by UV irradiation in flow. We supplemented the microgel precursor solution with glycidyl methacrylate (GMA) to introduce epoxide moieties into the microgel network, enabling us to specifically bind vitronectin (VTN) or other amine-bearing molecules to the microgels. Vitronectin was used to support iPSC adhesion and maintenance of the pluripotent state [45]. The incorporation of GMA was qualitatively analyzed by confocal imaging after post-functionalization with fluoresceinamine-isomer I (Fig. 1b). Here, specific binding was exclusively observed for GMA- modified microgels, revealing the successful incorporation of accessible epoxide functionalities into the microgel network.

**Fig. 1:**
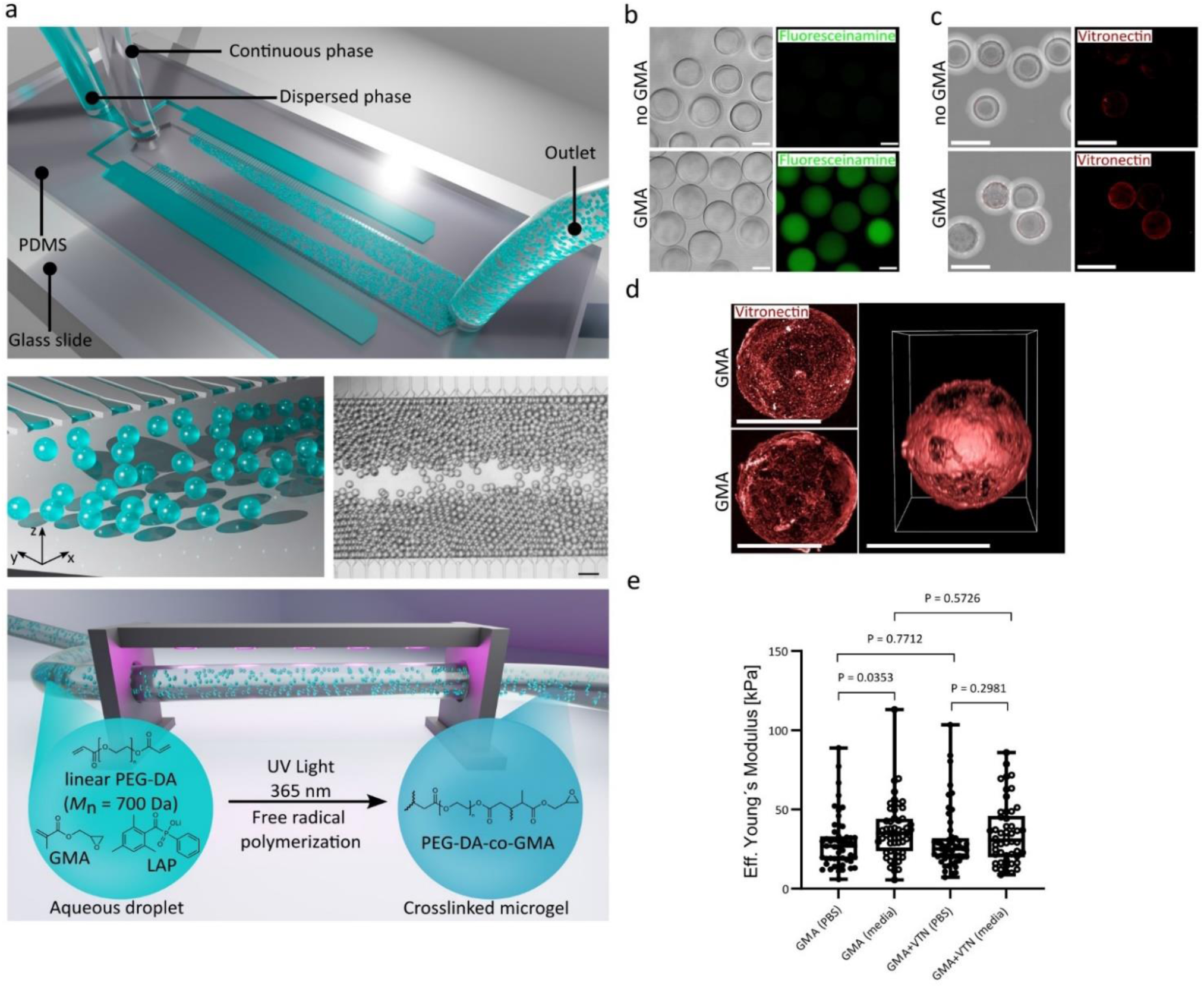
Large-scale production and functionalization of spherical microgels for iPSC-induced assembly. **a**, Top: Schematic of a parallelized step-emulsification microfluidic device used to produce spherical aqueous droplets, composed of PEG-DA, co-monomer GMA, and photoinitiator Lithium-Phenyl-2,4,6-trimethylbenzoylphosphinat (LAP). Middle: Schematic of droplet generation achieved via expansion of the aqueous phase from the shallow microchannel into the deep oil reservoir with corresponding microscope image (scale bar: 200 µm). Bottom: Schematic of in-flow UV polymerization in the outlet tube, transforming the pre-polymer droplets into polymerized PEG-DA-co-GMA microgels. Schematics are prepared with Blender. **b**, Confocal images of fluoresceinamine isomer I (green) labelled PEG microgels to demonstrate amine coupling to the GMA-modified microgels. Top: Pure PEG-DA microgels, revealed no specific binding of fluoresceinamine isomer I due to the lack of epoxy moieties. Bottom: PEG-DA-co-GMA microgels show specific binding of fluoresceinamine isomer I to the free epoxy moieties. Scale bars represent 40 µm. **c**, Confocal images of PEG microgels stained for VTN with fluorescent VTN-antibody. Top: PEG-DA microgels post-functionalized with VTN. Bottom: PEG-DA-co-GMA microgels post-functionalized with VTN. Scale bar represents 100 µm. **d**, Magnified confocal images of VTN functionalized PEG-DA-co-GMA microgels. Left: two distinct microgels exhibiting comparable VTN distribution on the surface. Right: 3D reconstruction visualizing the VTN distribution on a VTN functionalized PEG-DA-co-GMA microgel. Scale bars represent 50 µm. **e**, Mechanical properties of GMA modified and VTN post-functionalized microgels in different buffered solutions, determined via nanoindentation. Statistical analysis was performed via unpaired t-test (n=50-55).

Subsequently, we investigated the efficiency of VTN post-functionalization via qualitative confocal imaging after staining with a fluorescent VTN antibody (Fig. 1d). Sufficient coverage of VTN on the microgel surface was exclusively indicated for GMA-modified microgels. To investigate the mechanical properties of the microgels, we performed nanoindentation in different buffered solutions (Fig. 1e). Generally, stiffness was slightly lower in 1x PBS solution as compared to culture media, with no significant differences between GMA-modified microgels with or without VTN post-functionalization. For all tested conditions, the average stiffness ranges were below 50 kPa, which is within the relevant stiffness range for specific tissues *in vivo*, like the myocardium[46].

### Microgels enable mm-scale scaffolds for iPSC expansion and differentiation

Subsequently, biocompatibility and cell-material interaction with iPSCs were evaluated. The iPSCs adhere well on the VTN-functionalized microgels, which could be observed in the three-dimensional cell-microgel clustering (Fig. 2a). This demonstrated the necessity of both the GMA-incorporation and the VTN-functionalization, as microgel-cell cluster formation did not occur in conditions without the amine-epoxy-based biofunctionalization. Subsequently, we confirmed the preservation of pluripotency for the iPSCs cultured with the microgels for 4 days by staining for OCT4. Actin concentration around the microgels was observed (Fig. 2b), further indicating the proper iPSC-material interaction. With the confirmation of iPSC-material compatibility and the ability of the iPSCs to maintain their stemness, we developed a protocol for the formation of mm-scale microgel-iPSC constructs (Fig. 2c), subsequently referred to as scaffolds, which relies on the material-cell interaction to provide mechanical stability and integrity while allowing sufficient oxygen/nutrition diffusion.

**Fig. 2:**
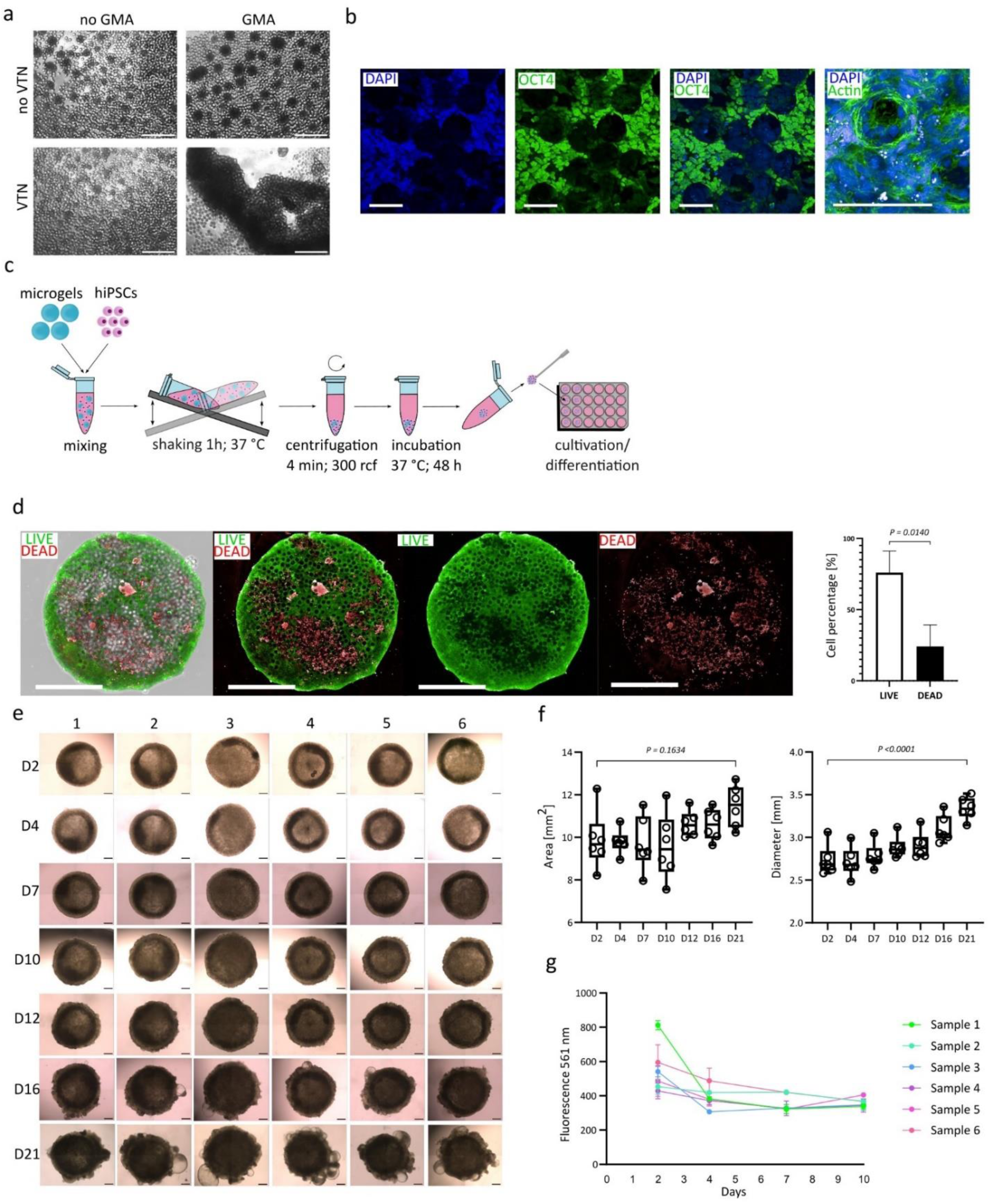
Production, viability, and reproducibility of 3D iPSC scaffolds. **a**, VTN functionalization via amine-epoxy coupling (GMA) to enable cell attachment to the microgels (brightfield images on day 4; scale bar is 500 µm). The incorporation of GMA and the functionalization with VTN are both essential to achieve adequate cell-material interactions. **b**, Cell material interaction between PEG-based microgels and iPSCs. Image 1-3: Pluripotency verification (OCT4) of iPSCs on PEG-DA-co-GMA microgels (DAPI (blue), OCT4 (green); scale bar is 100 µm). Immunofluorescence staining was performed on day 4. Image 4: Cell-material interaction demonstrated by Actin filament clustering at the interface. (DAPI (blue), Actin (phalloidin, green), scale bar is 100 µm). **c**, Schematic illustration of mm-scale scaffold production method (prepared with Inkscape) using iPSCs and PEG-microgels. **d**, LIVE/DEAD assay of scaffold after 2 days of culture (live cells (green), dead cells (red)). Right: Confocal Z-stack images (680 µm, Scale bar: 1 mm), Left: Cell viability analysis of LIVE/DEAD assay images using ImageJ (n=3, statistical significance was calculated via unpaired t-test). **e/f**, Reproducibility and shape controllability of scaffolds. Six samples (1-6) were analyzed over 21 days of culture (Scale bar: 500 µm). f, Area [mm^2^] and diameter (average of diameter in X- and Y-direction) [mm] were conducted with image analysis using ImageJ, n=6, statistical significance was calculated via One-way ANOVA and Tukeýs multiple comparison test. **g**, Metabolic activity assessment of 6 independent samples over a culture period of 10 days (alamarBlue assay, n=3 technical replicates per data point).

We optimized the iPSC-microgel ratio ∼400,000 cells/4,332 microgels (∼92 cells/microgel) (Supplementary Fig. 2), enabling reproducible generation of size- and shape-consistent scaffolds. These scaffolds were mechanically stable after 48 hours and could be transferred and cultured in solution in 24-well plates for the remaining culture or differentiation time. After 48 hours, we observed an average cell viability of 75.92 ± 15.21% (Fig. 2d), which is in a similar range as in conventional embryoid body formation, which often shows cell death of ∼ 10% after 48 hours and ∼ 50 % after 5 days[47]. The produced cell-biomaterial scaffolds had a mean area of 9.88 ± 1.35 mm^2^ (cross- section) and a diameter of 2.73 ± 0.17 mm after 48 hours of culture. Furthermore, we monitored the mechanical stability and reproducibility of the scaffolds demonstrating uniform spherical constructs that could be cultured in suspension without deformation or breakage over the course of 21 days (Fig. 2e). During this time, we observed continuous growth of the scaffolds up to an area of 11.47 ± 0.97 mm^2^ and a diameter of 3.34 ± 0.13 mm, demonstrating the iPSC expansion capability of the system (Fig. 2f). The metabolic activity was measured by the fluorescence shift caused through the enzymatic reduction of resazurin to resorufin by metabolically active cells and supported the reproducibility between multiple scaffolds. A high metabolic activity was measured after 48 hours, which balanced out to a stable level after 4 days (Fig. 2g). No necrotic core was formed as microgels provide stability and act as a “mock vasculature” facilitating oxygen, nutrient, and waste transport in the constructs, proposing a novel platform for the development of mm-scale organoids. Diffusion of oxygen through the cell/microgel constructs was simulated, suggesting a slight increase in oxygen diffusion due to the presence of microgels.

### Directed differentiation of microgel-iPSC scaffolds

We analyzed whether iPSCs maintained differentiation potential in the scaffolds by inducing differentiation toward the three germ layers. After 48 hours of expansion, we used directed differentiation protocols and analyzed the scaffolds after 7 days of differentiation on the protein, gene expression, and epigenetic level. Immunostaining for germ layer-specific markers (endoderm: GATA4; ectoderm: PAX6; mesoderm: Brachyury) showed expression of the respective markers upon directed differentiation (Fig. 3a). We further applied the PluripotencyScreen technology[1], which is based on a combination of site-specific DNA methylation (DNAm) assays to characterize iPSCs and their differentiation potential (Fig. 3b, c; Supplementary Fig. 4). A pluripotency score is indicative for pluripotency, whereas differentiation scores for mesoderm, endoderm, and ectoderm are indicative for early cell-fate decisions toward respective lineages (Fig. 3b). Deconvolution against reference matrix provides further estimates of the cellular composition (Fig. 3c). In the non-differentiated scaffolds, the iPSCs maintained a relatively high pluripotency score, albeit some DNA methylation changes point toward endodermal differentiation. This could be due to the high surface area presented by the microgels. Furthermore, three lineage differentiation evoked epigenetic modifications indicative of differentiation toward mesoderm, ectoderm, and endoderm, respectively (Fig. 3b). These findings were further supported by gene expression analysis of marker genes based on RT-qPCR (Fig. 3d, e). We thereby demonstrated the potential of iPSCs to assemble microgels into mm-sized constructs with the ability to expand until controlled differentiation is induced.

**Fig. 3:**
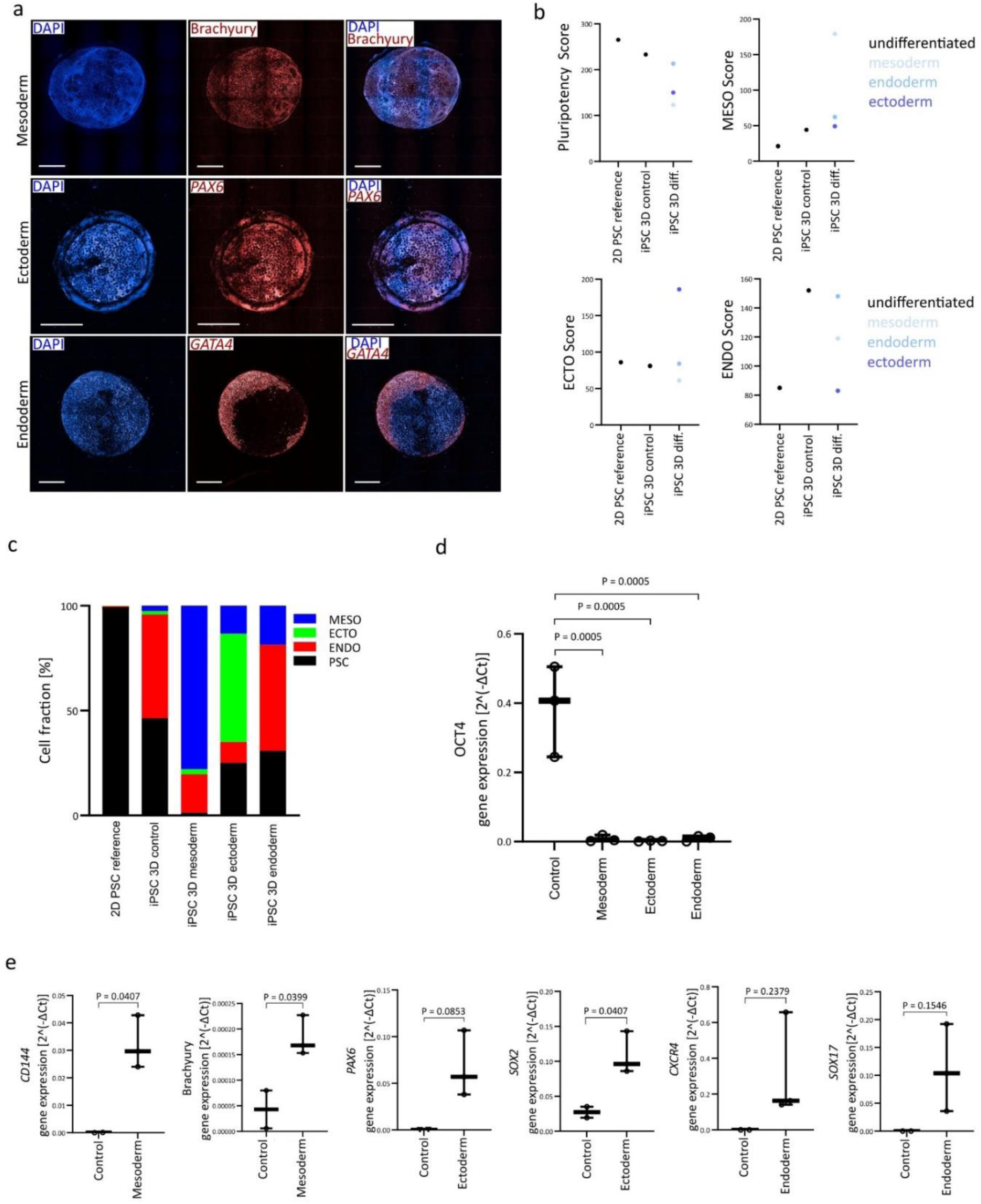
Differentiation potential of iPSC/microgel scaffolds into the three germ layers. **a**, Immunostaining of germ layer-specific markers after 7 days of directed differentiation (scale bar is 2 mm, DAPI (blue)). Endoderm (*GATA4* (red), confocal Z-stack images (666 µm)), ectoderm *(PAX6* (red), confocal Z-stack images (740 µm)), mesoderm (Brachyury (red), confocal Z-stack images (814 µm)). **b**, Pluripotency and differentiation scores inside scaffolds after 7 days of directed differentiation for three conditions: iPSC 2D reference (data from[1]), iPSC undifferentiated control (3D), iPSC differentiated samples (3D). **c**, Deconvolution of methylation data against reference matrix (data from[1]) demonstrating the different cell fractions in each sample. **d**, RT-qPCR of pluripotency marker OCT4 after 7 days of directed differentiation into each germ layer. Gene expression values were calculated using the 2^-ΔCt^- method, and statistical significance was calculated via One-way ANOVA and Tukeýs multiple comparison test (Control n=2, Sample: n=3; each n includes 4 pooled biological replicates.). **e**, RT-qPCR of germ layer-specific markers (endoderm*: CXCR4*, *SOX17*, ectoderm*: PAX6*, *SOX2*, mesoderm: *CD144*, Brachyury). Gene expression values were calculated using the 2^-ΔCt^- method, and statistical significance was calculated via unpaired t-tests (control n=2, sample n=3; each n includes 4 pooled biological replicates.).

Subsequently, we evaluated whether this technology can be used for cardiac organoid formation, as this reflects functionality and cell interconnectivity by spontaneous beating characteristics. After 48 hours of expansion within the microgel construct, the cardiac differentiation protocol was initiated. The scaffolds exhibited mechanical stability and shape-consistency during 23 days of cardiac differentiation (Fig. 4a). We further observed the reproducibility of shape and size of these organoids during the differentiation process, as well as the expression of the cardiomyocyte marker cardiac troponin T (cTnT) (Fig. 4b). The distribution of cTnT across the cardiac organoid demonstrated successful differentiation throughout the mm-sized organoid. The reduced cTnT signal observed in the center of the maximum projection confocal images (Fig. 4b) results from fluorescence intensity loss due to the samples’ density. While the edges can be imaged without significant attenuation across the entire Z-stack, the central regions experience signal reduction because of the limited laser penetration depth. Consequently, the maximum projection appears with higher fluorescence intensity at the edges compared to the center. The inside of the organoids was, therefore, analyzed with cryosections, displaying cell nuclei and cTnT expression, demonstrating the absence of a necrotic core (Fig. 4c). We validated cardiac differentiation with RT-qPCR of the marker genes *MHC-alpha*, *NKX2.5*, and *cTnT* at days 16 and 26 of differentiation. The 3D cardiac organoids demonstrated comparable marker gene expression to the two-dimensional control culture, disproving possible negative effects of the microgel-containing scaffold on cardiac differentiation. Furthermore, we observed a decrease in *MHC-alpha*[48] and *NKX2.5*[49] expression alongside an increase in *cTnT* expression[50] from day 16 to day 26 of differentiation, both suggesting the beginning maturation of cardiomyocytes (Fig. 4d). The cardiac organoids proved functional through consistent uniform beating, demonstrating their inherent pacemaker properties and the formation of an interconnected network throughout the whole scaffold (Supplementary Video 1,2). Calcium imaging within distinct regions of interest (ROIs) across the cardiac organoids exhibited equivalent beating frequencies, validating the cardiomyocyte interconnectivity without hindrance by the microgels (Fig. 4e). These results suggest the successful formation of mm-sized cardiac organoids using iPSCs-microgels-based scaffolds.

**Fig. 4:**
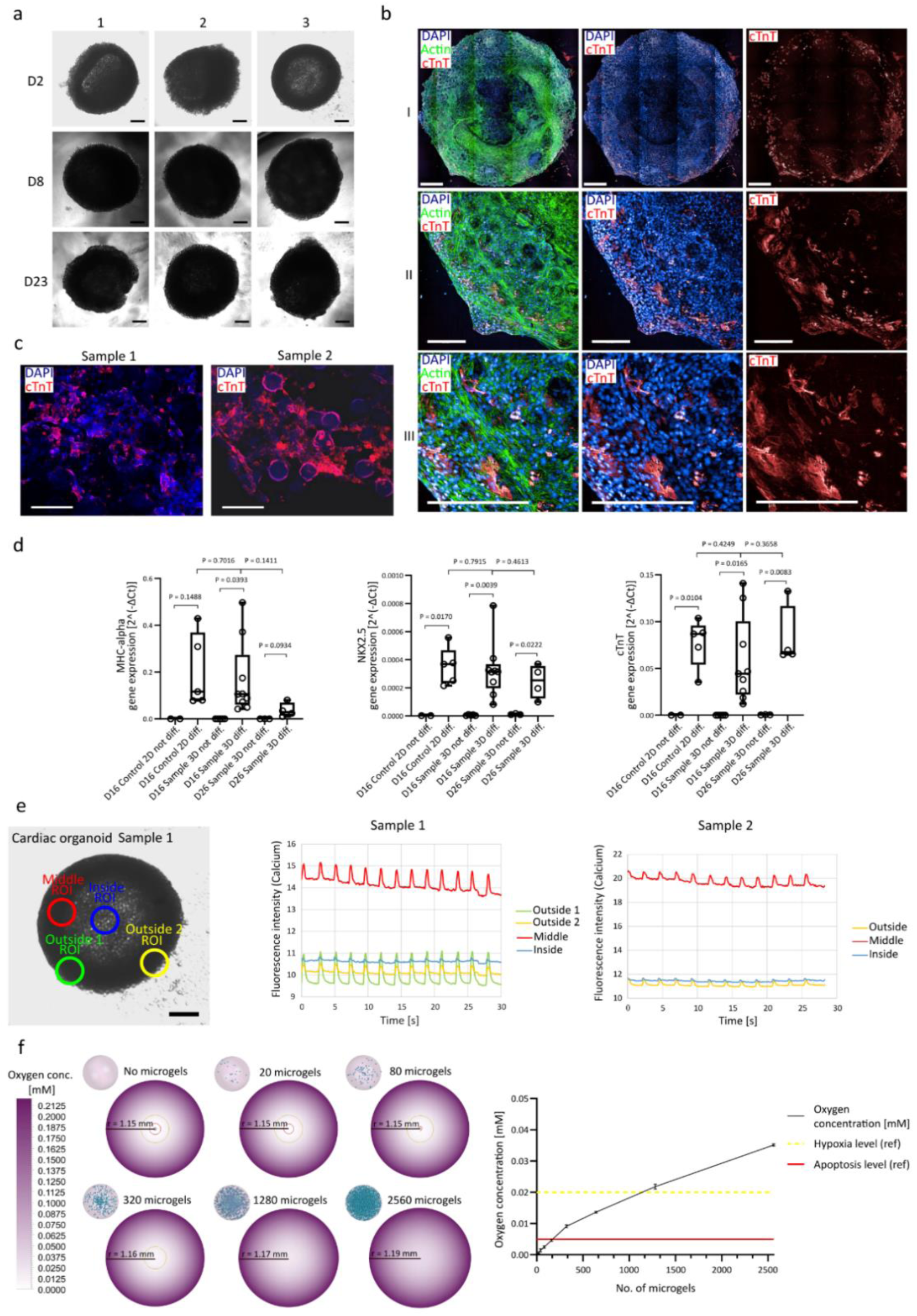
Differentiation into cardiac organoids. **a**, Uniform shape of cardiac organoids over 23 days of culture (3 different samples (1-3), scale bar is 1 mm). **b**, Immunofluorescence confocal imaging of cardiac organoids on day 23 of culture (DAPI (blue), Actin (Phalloidin, green), cTnT (red)). Confocal Z-stack images (720 µm), scale bar 1 is 1 mm, 2 and 3 are 200 µm). **c**, Cryosections from the middle (Z-axis) of cardiac organoids on day 23. Cryosections were stained for DAPI (blue) and cTnT (red), demonstrating cardiac differentiation inside the construct. **d**, RT-qPCR of cardiac-specific markers (*MHC-alpha*, *NKX2.5*, *cTnT*) on day 16 and day 26 of differentiation with respective undifferentiated controls and a 2D comparison. Gene expression values were calculated using the 2^-ΔCt^- method, and statistical significance was calculated via unpaired t-tests (n=3 – n=9; each n includes 4 pooled biological replicates.). **e**, Calcium imaging of cardiac organoids (performed at one Z-plane). The different ROIs (outside, middle, inside) were set and analyzed with NIS Elements software demonstrating consistent cardiomyocyte contraction frequency (fluorescence intensity changes) at respective ROIs. Left image: general positioning of ROIs respective to cardiac organoid (Sample 1) (scale bar is 500 µm). Right: graphs representing calcium imaging data in each of the ROIs for two samples. **f**, Simulation results of oxygen transport. Left: contour plots of oxygen concentration. Simulation of oxygen transport in a cardiac organoid with a 1.15 mm radius shows the formation of a necrotic core (C_Oxygen_ < 0.005 mM) and hypoxic area (0.005 mM < C_Oxygen_ < 0.02 mM). Adding microgels to the organoid enhances oxygen transport impeding necrotic core formation at more than 1280 microgels per organoid. Right: correlation between minimal oxygen concentration and microgel volume ratio determined through simulation. Further details regarding oxygen transport modeling and simulation are reported in supplementary.

To further support our hypothesis, we propose that microgels enhance oxygen diffusion within scaffolds and help prevent the formation of a necrotic core. To test this, we simulated oxygen diffusion inside cardiomyocyte spheroids containing 600,000 cells, which is estimated to be in a comparable scope as the experiment after proliferation. The oxygen diffusion within the microgel-supported scaffold was investigated through numerical simulations based on the transport diffusion equation. The steady-state oxygen concentration profiles were obtained by solving the governing partial differential equation using a finite volume method. The simulation demonstrated an increase in oxygen concentration throughout the scaffold with increasing microgel amount (Fig. 4f). With approximately 1,280 microgels added, the formation of a necrotic core (C_Oxygen_ < 0.005 mM[51]) and hypoxic area (0.005 mM < C_Oxygen_ < 0.02 mM[52]) could theoretically be circumvented. Thereby, the simulation supports our observation that with an even higher number of microgels (4,335 microgels/scaffold), we could generate mm-sized cardiac organoids without necrotic core-formation.

### Scale-down of scaffold production to 96-well format and differentiation to cardiac organoids and retinal photoreceptor cells

We scaled down the proposed microgel platform for iPSC expansion and differentiation to a 96-well format, allowing for smaller scaffold production with higher throughput. U-shaped well plates were employed without the necessity of centrifugation, simplifying the process and increasing applicability (Fig. 5a). After 48 hours of iPSC expansion inside the microgel-based scaffolds (337 cells/microgel), the cells were differentiated into stable cardiac organoids. This cell/microgel ratio is higher than for the mm-scale organoids, likely due to the fact that no centrifugation step is included. The cross-section of the organoids had an area of 0.96 ± 0.13 mm^2^ (Supplementary Fig. 6), which is a reduction in size of approximately 3.5-fold compared to the millimeter-size scaffolds (Fig. 2f). Figure 5b shows a single microgel as a representative view of the uniform cell monolayer covering the microgel surface, indicating a strong cell-material interaction. Cardiac differentiation was confirmed via immunofluorescence staining (cTnT) throughout the whole scaffold and uniform beating across the organoids (Fig. 5c; Supplementary Video 4,5). The cTnT fibers are directly adjacent to the microgels arranged around the microgel surface (Fig. 5d), and generate characteristic striations, implying that the microgels support the formation of a functional and interconnected cTnT network (Fig. 5e).

**Fig. 5:**
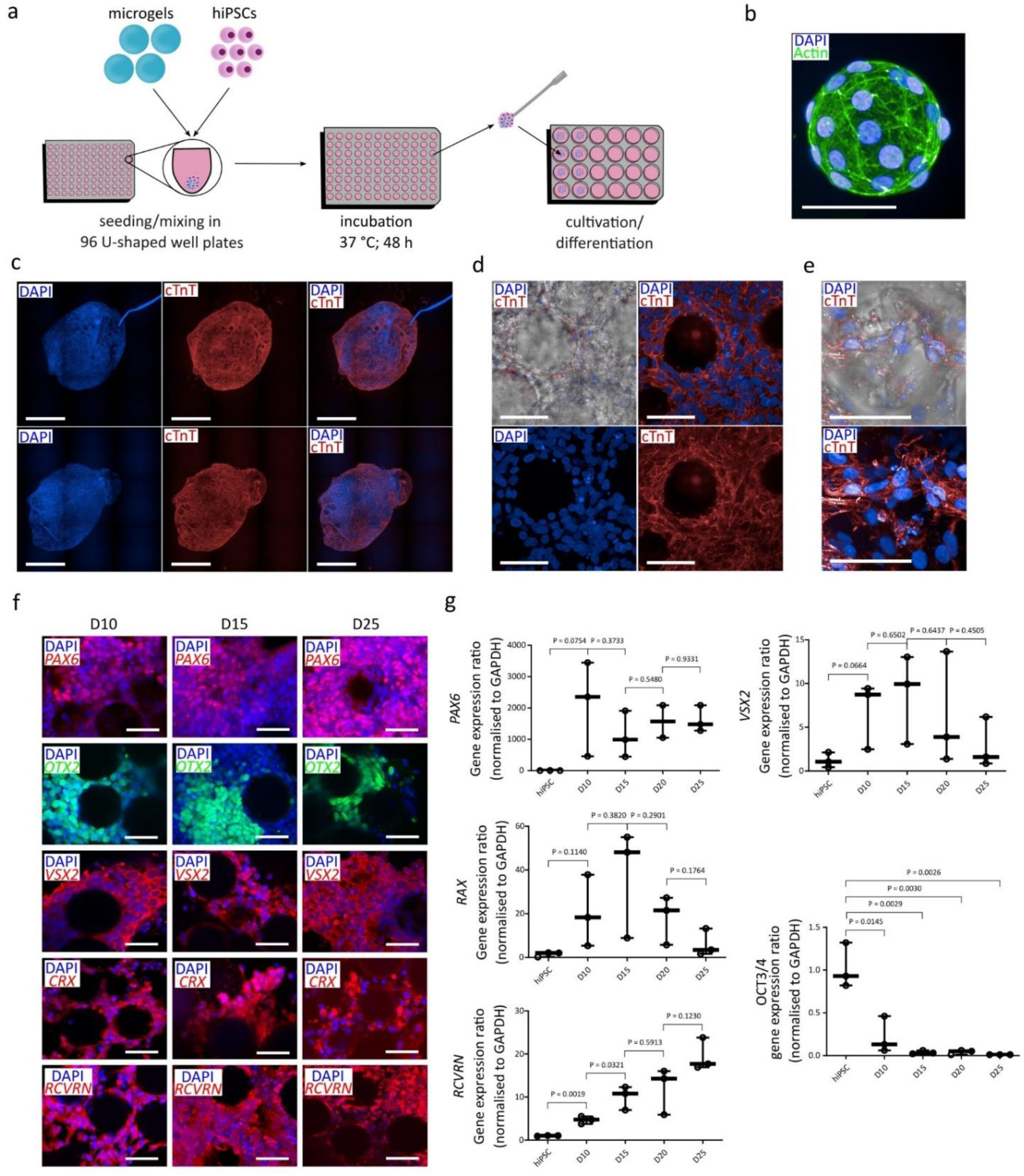
Scale-down of scaffold production and differentiation. **a**, Schematic illustration of iPSCs/microgels scaffold production in a 96-well plate (prepared with Inkscape). **b**, Single microgel with a cell monolayer broken of the scaffold during staining after 30 days stained for Actin (green) and DAPI (blue). Scale bar is 50 µm. **c**, Immunofluorescence confocal imaging of cardiac organoids on day 30 of culture stained with DAPI (blue) and for cTnT (red). Confocal Z-stack images (540 µm), scale bar is 500 µm. **d**, Immunofluorescence images of microgels embedded in cardiac organoid. Confocal Z-stack images (30 µm) and the scale bar is 50 µm. **e**, Immunofluorescence images of distinct cTnT structure in cardiomyocytes surrounding a microgel inside a cardiac organoid. Confocal Z-stack images (200 µm) and the scale bar is 50 µm. **f**, Immunofluorescence images of retinal differentiation of scaffold with distinct retinal markers (PAX6 (red), OTX2 (green), VSX2 (red), CRX (red), RCVRN (red)) on day 10, 15, and 25 of culture. Confocal Z-stack images (xx µm) and the scale bar is 50 µm. **g**, RT-qPCR of retinal differentiation (*PAX6*, *VSX2*, *RAX*, *RCVRN*) and pluripotency markers (*OCT3/4*). Gene expression ratios were calculated with a script provided by TopTipBio and statistical significance was calculated via unpaired t-tests (n=3).

As a demonstration of the applicability of this scaffold approach for different tissues, we also differentiated the constructs into retinal photoreceptor cells. iPSC-based microgel scaffolds (185 cells/microgel) were assembled, and we initiated the differentiation by adapting a 2D photoreceptor differentiation protocol to our 3D system[53]. The cell/microgel ratio here differs from the one applied for cardiac organoids, since a higher initial cell number was used for retinal photoreceptor differentiation. With higher cell numbers, the proliferation-related increase of cells is higher than with lower starting cell numbers, while the microgel amount stays constant. We observed thereby, that the lower the initial cell number used, the higher the cell/microgel ratio should be for the formation of stable scaffolds within 48 hours using this seeding method. Throughout the differentiation process, we observed key milestones in photoreceptor development. Retinal cells were characterized at various stages by immunofluorescence microscopy and qRT-PCR (Fig. 5f, g). By day 10, there was a notable increase in the expression of eye field transcription factors *PAX6* and *RAX* compared to undifferentiated iPSCs. Furthermore, we detected *VSX2*, a marker specific to retinal progenitor cells, and *CRX*, a marker of photoreceptor progenitors. *CRX* expression first observed in the microgel scaffold at day 10 occurs earlier than reported in retinal organoids and photoreceptor differentiation protocols from iPSCs, which were described to take at least 21 days.[53–56]. This earlier differentiation is likely due to the 3D system, which provides a more conducive environment, along with the physiological stiffness of the microgels compared to tissue culture plastic. The expression of recoverin, a calcium-binding protein crucial for phototransduction, increased consistently over the course of 25 days.

Using the U-shaped 96-well plates, we demonstrated successful differentiation of the expanded iPSCs inside the microgel scaffolds into two different targets, illustrating the applicability of the proposed technology for variable use in organoid and stem cell research, with advantages such as necrotic core prevention or faster differentiation.

### High-throughput production of iPSCs/microgels scaffolds using an automated liquid handling system

After establishing the iPSC-microgel technology for the 96-well plate format, the next objective was to apply the proposed system in an automated liquid handling system for high-throughput automated production of 3D constructs with potential application for compound screening. We developed a method to enable the formation of iPSC/microgel scaffolds for organoid formation in low-volume (25 µL) flat surface 384-well plates (Fig. 6a). The scaffold formation in this manner is solely driven by the self-assembly of the iPSCs with the microgels into 3D spherical constructs. This cell-driven assembly was highly dependent on the cell-to-microgel ratio, which was optimized to allow for the formation of one single scaffold per well (Supplementary Videos 6-10). Supplementary Fig. 7a demonstrates how iPSCs use microgels at different cell-to-microgel ratios to assemble into sizable spherical structures in a fully automated workflow. Based on stability and shape of the iPSC/microgel scaffolds, a ratio of 125 cells/microgel was selected with 10,000 iPSCs and 80 microgels (Supplementary Fig. 7a). We observed good scaffold formation and the lowest number of dead cells or non-incorporated microgels for this condition. The increase in the cell number to more than 20.000 per sample caused an increase in cell debris, indicating cell death of non-incorporated iPSCs. For this ratio, we observed the formation of spherical scaffolds after 24 hours, with a subsequent stabilizing and densifying of the construct within 48 hours, resulting in an average area of 0.19±0.05 mm^2^. The size of the scaffolds increased consistently up to an area of 0.25 ± 0.03 mm^2^ after 5 days of culture, pointing out iPSC expansion in a high cell-density scaffold (Fig. 6e, f). Additionally, the possibility of scaffold transfer through automated pipetting was demonstrated, allowing for the removal of cell debris during culture time (Supplementary Fig. 7), needed for an optimized culture workflow. This last step is essential for the automated handling of the organoids during long-term culture. We observed microgels incorporated and distributed throughout the scaffold (Fig. 6d). Furthermore, we demonstrated the applicability of this method for a large number of samples (160) in an automated manner, depicting the potential for large-scale organoid production or drug screenings in the future (Supplementary Fig. 8).

**Fig. 6:**
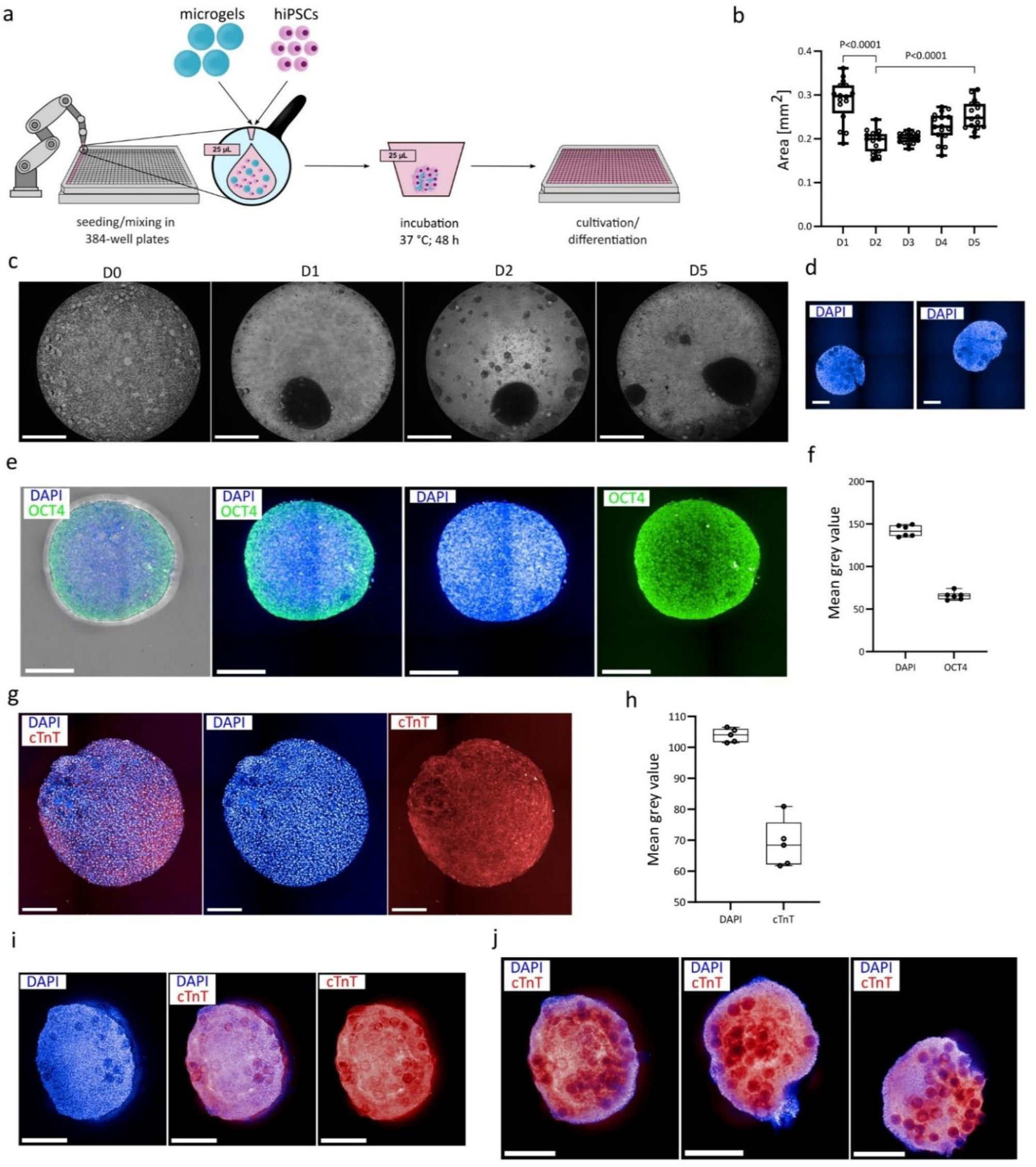
Automated production of scaffolds. **a**, Schematic illustration of iPSCs/microgels scaffold production method inside 384-well plates (prepared with Inkscape) using an automated liquid handling system. **b**, Scaffold area analysis over 5 days of culture. Compaction of scaffolds from first to second day with subsequent scaffold growth. Statistical significance was calculated via One-way ANOVA and Tukeýs multiple comparison test (n=16). **c**, Scaffold formation and culture in low-volume 384-well plates at day 0, 1, 2, and 5 (scale bar: 500 µm). **d**, Middle view of the scaffold after 2 days of culture (DAPI (blue)). Scale bar: 200 µm. Left image: segment (37 µm) from the middle of the Z-stack image (324 µm). Right image: segment (18 µm) from the middle of the Z-stack image (333 µm). **e**, Pluripotency of scaffolds after 2 days (DAPI (blue), OCT4 (green)). Scale bar: 200 µm. Confocal Z-stack (288 µm). Mean grey values of DAPI and OCT4 stains in scaffolds at day 2. Low standard deviation demonstrated reproducible cell number and pluripotency for automated scaffold production. **f,** Mean grey values of DAPI and OCT4 stains in scaffolds at day 2. Low standard deviation demonstrated reproducible cell number and pluripotency for automated scaffold production (n=6). **g**, Cardiac differentiation of scaffolds day 30 (DAPI (blue), cardiac Troponin T (cTnT) (red)); Confocal Z-stack images 10x magnifaction (296 µm) (scale bar: 200 µm). **h**, Mean grey values of DAPI and cTnT stains in scaffolds at day 30. Low standard deviation demonstrated reproducible cell number and differetiation for automated scaffold production (n=5). **i**, Cardiac differentiation of scaffolds day 30, a clearing protocol was used to visualize microgels inside of the constructs; (DAPI (blue), cardiac Troponin T (cTnT) (red)); Confocal Z-stack images 10x magnifaction (296 µm) (scale bar: 200 µm). **j**, Single plane at 59.2 µm depth of cardiac differentiated scaffolds for the demonstration of microgel distribution throughout the construct and uniform differentiation. (DAPI (blue), cardiac Troponin T (cTnT) (red)); Confocal Z-stack images 10x magnifaction (296 µm) (scale bar: 200 µm).

To validate the compatibility of the high-throughput system for iPSC-based applications, the pluripotency of the automatically produced scaffolds was confirmed after 48 hours by immunostaining for OCT4 (Fig. 6e). Mean grey value analysis of the confocal DAPI and OCT4 stained images exhibited low standard deviations of 4.6 % (DAPI) and 7.4 % (OCT4), indicating reproducible cell numbers of pluripotent cells (Fig. 6e). Furthermore, OCT4 expression, as an indicator for pluripotent state, was similar in cells directly adjacent to the microgels with cell-material contact as compared to cells further away from a microgel, only experiencing cell-cell contacts. We thereby conclude that iPSCs and microgels, processed by the automated pipetting system, also form 3D constructs with iPSCs maintaining pluripotency. To demonstrate the applicability of the high-throughput system for organoid production, we generated cardiac organoids by inducing cardiac differentiation after 48 h of scaffold formation under iPSC maintenance conditions. After 30 days of differentiation, we observed spontaneously beating (Supplementary Video 11) cardiac organoids, which furthermore expressed cardiac Troponin T (cTnT) homogeneously throughout the whole scaffold (Figure 6g, i). The mean grey value analysis of DAPI and cTnT expression in these organoids demonstrated reproducibility of the production and a reproducible differentiation (Figure 6h). Figure 6j additionally demonstrates the uniform differentiation of cardiac organoids without necrotic core formation or reduced differentiation towards the center of the organoid. These results of organoids produced with an automated liquid handling system demonstrated the applicability of the iPSC/microgel scaffolds for high-throughput production to function as a platform technology for organoid generation in an automated, reproducible manner, which could be used for drug discovery.

## Conclusion

In stem cell and organoid research, challenges regarding reproducibility and scalability prevail, complicating the reliable application of these technologies for drug development and disease modeling. To tackle some of these challenges, we have developed a new technology that combines iPSC expansion and differentiation within the same construct and facilitates the reproducible manufacturing of organoids from sizable to high-throughput. This approach is entirely chemically defined, employing PEG-based microgels, coated with recombinantly produced vitronectin, as the matrix material, and using xeno-free media. We believe that this technology can provide a user-friendly foundation for advancing chemically defined organoid and stem cell research for many different tissue models with a significant impact on tissue engineering and pharmacology. Application of more specialized microgel systems, which allow for introducing features such as larger pore sizes[40, 41], mechanical actuation[59], and alignment[60] for controlling tissue architecture and differentiation, is among the interesting opportunities to be embedded into this technology. Additionally, the incorporation of the microgels could be further developed to allow for morphogen release[61] or structural gradient formation[62], proposing a potential for the generation of spatially defined compartments within one organoid. These advancements could open up a wide range of new possibilities for organoid development and maturation, enabling more relevant models that could better recapitulate native tissue architecture and lead to more mature organoids. The scalability and reproducibility of the method could furthermore aid in long-term cultivation[63] and complex patterning[62], contributing to advancements in disease modelling and high-throughput drug screening. Other methods proposed to allow for long-term cultivation or mm-scale organoids are currently not automation-compatible, since they require, for example, organoid slicing or stirring cultures to allow for the needed nutrient and oxygen supply[64, 65]. Furthermore, the platform demonstrates adaptability to different tissue types, proposing a potential in supporting structural and functional aspects across diverse biomedical applications. In the long term, we believe this platform could serve as a modular foundation for building complex, mature, functional tissue systems suitable for translational research. Future research must further tune these materials for specific tissue types, which requires close collaboration between stem cell researchers and materials scientists. The insights gained from this research could bring organoid studies closer to reducing or replacing animal testing, while also providing human-based alternatives to improve and accelerate drug development and personalized medicine in the future.

## Materials and methods

### Microfluidic Master Mold Fabrication

A positive 3D structure of the microfluidic channel geometry was designed (AutoCAD 2019), processed in DeScribe 2.7, and was fabricated using two-photon direct laser writing (Nanoscribe GmbH, Photonic Professional GT). As previously described[66], an acryl-silanized glass coverslip (LABSOLUTE® Th. Geyer 76 x 26 x 1 mm) was covered with a photoresist (Nanoscribe, IP-S) and mounted in the 3D printer. The print was performed with a 25x objective (Blocksize X: 200 Y: 200 Z: 160 µm, Shear 15°). After printing completion, the master structure was submerged in propylene glycol monomethyl ether acetate (PGMEA, Sigma-Aldrich, ≥ 99.5%) for 10 min to remove uncrosslinked polymer resin, followed by washing with isopropanol (IPA) solution for 3 min. The substrate was dried under nitrogen flow and post-cured for 4-6 hours using UV light (200 mW, λ=365 nm). The cured structure was fluoro-silanized with trichloro-(1H,1H,2H,2H-perfluoroctyl)- silane (50 µL per glass coverslip) in a desiccator under high vacuum overnight.

### Microfluidic Chip Fabrication

A mixture of PDMS base and curing agent (10:1 w/w, Sylgard® 184, Dow Corning) was cast on the master structure, degassed, and cured at 60 °C for 12 hours. The PDMS replica was cut from the master mold, inlet holes were punched (0.75 mm biopsy puncher, Electron Microscopy Science), and cleaned with IPA and Milli-Q water. The glass slide (76 x 52 x 1 mm, Marienfeld, Germany) was prepared analogously, including additional washing steps with acetone. The PDMS slab and glass slide were plasma treated (PVA TePla 100 Plasma System, 28 mL/min O_2_, 100 W, 40 s) and bonded together. The microfluidic chip was stored at 60 °C for 12h to complete the bonding. Before use, the device was rendered hydrophobic by chemical vapor deposition (CVD) with trichloro-(1H,1H,2H,2H-perfluoroctyl)-silane (50 µL per chip) in a vacuum desiccator overnight. Afterward, the outside of the chip was rinsed with Novec™ 7500 (3M) and wiped using lint-free tissues.

### Microgel Production

Epoxy-functionalized microgels were prepared from aqueous solutions composed of polyethylene glycol diacrylate (PEG-DA, 15% w/w, Mn 700, Sigma-Aldrich), Lithium-Phenyl-2,4,6-trimethylbenzoylphosphinate (LAP, 1% w/w, Sigma-Aldrich) photoinitiator, and freshly distilled glycidyl methacrylate (GMA, 1% w/w, Sigma-Aldrich). As a control, unfunctionalized PEG microgels were produced analogously without the addition of GMA. The solution was stored in brown glass vials until use to protect from undesired light exposure. The continuous oil phase was composed of Novec™ HFE 7500 with Krytox™ 157 FSH surfactant (4.5% v/v, DuPont). Both solutions were transferred into a syringe (Hamilton Gastight® Series 1000, 10mL), and the syringe containing the dispersed phase was wrapped in aluminum foil. A 25G cannula was inserted into PTFE tubing (OD 0.9 mm, ID 0.4 mm, TECHLAB GmbH) and attached to the syringes. Residual air was removed, and the syringes were fixed in a syringe pump (Havard Apparatus Pump 11 Elite). The other ends of the tubes, as well as an outlet tube (Polyethylene, OD 0.038 in ID 0.023 in, Instech Laboratories), were inserted into the dedicated inlet holes on the microfluidic device. The continuous phase was started at a flow rate of 1000 µL/h to prefill the collection reservoir. Afterwards, the dispersed phase was started at 1000 µL/h until droplet formation was visible. The flow rates were then adjusted to 3000 µL/h and 2000 µL/h for the continuous and dispersed phase, respectively. Droplet crosslinking was achieved by guiding the outlet tube through a self-constructed UV-LED set-up, as previously reported[67]. Briefly, the UV set-up contains a fixed frame featuring an array of five LEDs (Starboard, Luminus SST-10-UV-A130, 365 nm) spaced at a distance of 23 mm, powered by a 24 V power supply, and operating at an irradiation dose of 30.36 mW/cm^2^. The microgels are crosslinked in-flow by fixing the outlet tube in a PDMS gasket connected to the frame. Purification of the microgels was achieved by multiple subsequent washing steps, as previously reported.[68] In short, the excess continuous phase was pipetted off and the microgels were sequentially washed with Novec™ HFE 7500, n-hexane + span 80 (1% v/v), n-hexane, IPA, and Milli-Q water. Each solvent was applied five times before proceeding to the next solvent. After the purification, the microgels were stored in Milli-Q water at 4 °C until use.

### Epoxide labeling with fluoresceinamine-isomer I and qualitative confocal imaging

Incorporation of GMA was qualitatively analyzed by amine-epoxide coupling with fluoresceinamine-isomer I. Fluoresceinamine in DMSO (5 mg/ml, 100 µL) was added in excess to a dispersion of PEG-DA-co-GMA and PEG-DA microgels in water (30 µL jammed microgel sediment, 900 µL Milli-Q water), yielding a final fluoresceinamine concentration of 0.5 mg/ml. Amine-epoxy coupling was achieved overnight on a roller. The microgels were then washed twice with DMSO and six times with Milli-Q water to remove excess of unbound fluoresceinamine. Confocal laser scanning microscopy was performed on a Leica TCS SP8 microscope (Leica Microsystems, Germany). Fluoresceinamine was excited at 𝜆 = 488 nm via an argon laser. The HC PL FLUOTAR 10x/0.30 DRY objective was used to detect the resulting emission at 𝜆 = 500– 555 nm.

### Nanoindentation

The mechanical properties of the microgel particles were determined using a Pavone Nanoindenter (Optics11Life, Amsterdam, The Netherlands) equipped with a cantilever-based analysis probe with a spherical tip radius of 10 µm and a cantilever stiffness of 0.48 N/m. Prior to measurement, 40 µL of microgel dispersion was pipetted into a 24-well plate, followed by the addition of 1 mL buffered solution, as indicated. To ensure attachment of the microgels to the well plate surfaces, the well plate bottom was pre-treated overnight with a poly-L-lysine solution (50 µL, 0.1% w/v in water, Sigma-Aldrich). All measurements were performed at room temperature. The microgel particles were indented p to a maximum load of 1.2 µN and a piezo speed of 1 µm/s. The obtained load-indentation curves were used to calculate the effective Young’s modulus using the Hertzian contact model, fitted to a maximum indentation depth of 1600 nm. The data analysis was performed with Dataviewer V 2.5 (Optics11Life).

### Vitronectin-functionalization of microgels

Prior to functionalization with vitronectin, the microgels, suspended in 1x PBS pH 7.4, were UV-sterilized for 2 hours and subsequently handled only under sterile conditions. Microgels were functionalized with the Vitronectin XF™ solution (STEMCELL technologies^TM^). Therefore, microgels were pelleted and resuspended in an equal volume of 1xPBS corresponding to the pellet volume (e.g., 100 µL microgel pellet + 100 µL 1x PBS). To this, equal volumes of the VTN-solution (e.g., 100 µL) in a concentration of 250 µg/mL were added, reaching a final concentration of 125 µg/mL. The final solution was incubated overnight at 4 °C and for 20 minutes at 37 °C directly prior to use for experiments.

### Cell culture

Two iPSC lines (UKAi009-A and UKAi0011-A) were used in this study, and further information can be found in the Human Pluripotent Stem Cell Registry (hPSCreg)[69]. Maintenance cell culture of iPSCs was performed in tissue culture-treated 6-well plates coated with Vitronectin (Vitronectin XF^TM^ STEMCELL technologies^TM^,# 07180; 0.5 µg/cm^2^) in StemMACS™ iPS-Brew XF (Miltenyi biotec, 130-104-368). Cells used during this work were between passages 18 and 53.

#### Differentiation into three germ layers

Germlayer differentiation was performed on scaffolds on day 2 using the StemMACS™ Trilineage Differentiation Kit (Miltenyi biotec, 130-115-660), yielding fully differentiated samples after 7 days (endoderm 5 days of differentiation) of differentiation according to the supplier protocol. After 7 days of differentiation (9 days of culture), samples were further analyzed by RT-qPCR, epigenetic analysis, or immunofluorescence staining.

#### Differentiation into cardiac organoids

Cardiac organoid differentiation was performed starting after 48 hours of scaffold formation using the StemMACS™ CardioDiff Kit XF (Miltenyi biotec, 130-125-289). Media was prepared as described in the protocol. Time points of mesoderm and cardiac induction were applied as described in the protocol, while media change was performed daily for the whole culture time, differing from the protocol provided by the supplier. Cardiac organoid culture was performed for different culture periods ranging from 16 to 31 days. Samples were further analyzed by RT-qPCR and immunofluorescence staining after culture.

#### Differentiation of iPSCs to photoreceptor progenitors

Human induced pluripotent stem cells (iPSC-106) were cultured on vitronectin (STEMCELL technologies)- coated tissue culture plate. After reaching 70-80% confluency, the iPSCs were seeded with microgels in a 96- well ultra-low attachment U-bottom plate and maintained in iPS-Brew medium (Miltenyi Biotec) for 48 hours to promote embryoid body (EB) formation. Retinal differentiation was performed based on a published protocol with suitable modifications [53]. Unlike the published protocol, EBs were maintained in suspension culture throughout differentiation rather than plating them onto Matrigel-coated surfaces. Retinal induction was initiated by replacing the iPS-Brew medium with Retinal Induction Medium (RIM), which consisted of DMEM/F12 (Gibco), 10% KSR (Gibco), 0.1 mM non-essential amino acids (Sigma), 1% N2 (Gibco), 2% B27 without vitamin A (Gibco), supplemented with 10 ng/mL Noggin (STEMCELL Technologies), 10 ng/mL DKK 1 (STEMCELL Technologies), 10 ng/mL IGF-1 (R&D Systems), and 5 ng/mL basic fibroblast growth factor (bFGF; Millipore). After 5 days, the medium was replaced with RIM without KSR, and the cultures were maintained under 3D suspension conditions until day 25.

### Assessment of vitronectin-functionalization

VTN-functionalized microgels (description above) were pelleted, and 25 µL of pellet volume was pipetted into a 96-well plate. Afterward, 200.000 iPSCs (1000 cells/µL) were added to each well, and iPS-Brew was used as the media. For the first 24 hours, 10 µM of ROCK-inhibitor (Y-27632 (Dihydrochloride), STEMCELL technologies^TM^, # 72304) was added. Media was exchanged daily, and after 4 days of culture, the cell-material interaction was assessed via brightfield microscopy.

### Microgel-iPSC scaffold formation

After reaching around 80% confluency, iPSCs were detached into single cells using ACCUTASE^TM^ (STEMCELL technologies^TM^, # 07920) cell detachment solution. Afterwards, microgels stored in the coating solution were centrifuged (1 min, RT, 8000 rpm) and the coating solution was replaced with iPS-Brew media, in a ratio of 4:1 media to microgel pellet volume (e.g., 400 µL media + 100 µL microgel pellet). The respective microgel number added to the experiments using this microgel solution was calculated using a calibration curve (Suppl. Fig. XX). To all prepared experiments, ROCK-inhibitor (Y-27632 (Dihydrochloride), STEMCELL technologies^TM^, # 72304) was added in a concentration of 10 µM for the first 24 hours.

#### Mm-scale scaffolds

For the preparation of mm-scale scaffolds, 100 µL microgel solution (explained above) and 400.000 cells (1000 cells/µL) were added to a 1.5 mL protein LoBind Eppendorf^TM^ tube and filled up with iPS-Brew media to a volume of 800 µL. Afterwards, the mixture was put on an orbital shaker for 1 hour inside the incubator (37 °C, 5% CO_2_), to allow for initial attachment of the cells to the microgels. Subsequently, the mixture was centrifuged for 4 min at 300 g (RT), and the resulting pellets were cultured for 48 hours inside the Eppendorf ^TM^ tubes in the incubator with 700 µL media exchange after 24 hours. On the second day, formed scaffolds were removed from the tubes using a small spatula and transferred into 24-well plates filled with 500 µL media per well. They were either cultured in iPS-Brew media or differentiated (see below) with daily media change.

#### 96-well plate scaffold production

Either 6.25 µL (maintenance and cardiac differentiation) or 25 µL (retinal differentiation) microgel solution was pipetted into Nunclon™ Sphera™ U-well 96-well plates (Thermofisher scientific^TM^, 174925) with 100.000 cells (maintenance and cardiac differentiation) or 200.000 cells (retinal differentiation), respectively. The volume was filled up to 250 µL per well, and 200 µL of iPS-Brew media was changed after 24 hours. On the second day, the formed scaffolds were transferred into 24/48-well plates using a small spatula and cultured either in maintenance or differentiation media (see below) with daily media change (500 µL).

#### Scaffold production in 384-well plate with automated liquid handling system

JANUS G3 automated liquid handling workstations (Revvity, USA) was used for automated production of scaffolds in high-throughput format. All aspiration, dispensing, and mixing procedures were performed using Varispan dispensing arm. A 384 low-volume well plate (Greiner Bio-One, Netherlands) was used. Initially, the above-described microgel solution and a cell solution with a concentration of 5×10^6^ cells/mL were prepared in two separate 1.5 mL Eppendorf^TM^ tubes (protein low-bind in case of microgels), which were used with the automated liquid handling system to prepare a master mix of cells and microgels in a third Eppendorf tube. Each well was filled with 5.25 µL of cell-microgel master mix (1.25 µL microgel solution, 4 µL cell solution) and 19.75 µL iPS-Brew media. To minimize the evaporation from experimental wells, adjacent wells to the experimental ones were filled up with 25 µL 1x PBS. After one day of culture, the cells and microgels self-assembled into an organoid-like scaffold lying on the bottom of the well plate. Then, the daily media change was performed by discarding 17 µL of old media from the top of the well using the liquid tracking option. The wells were then filled with the same volume of new culture media. Note that each step requires proper aspiration and/or dispensing vertical position with respect to the bottom of the wells and proper liquid tracking setup to ensure precise dispensing/aspiration, and avoid cross-contamination and bubble formation. Additionally, we were able to transfer the organoid from well to well, hence removing cell debris by gently aspirating 15 µL at 1 mm above the well bottom and dispensing it in a new well using an automated liquid handling system.

#### Immunostaining and fixation of scaffolds

Scaffolds were fixed with 4 % PFA for 45 minutes and afterward washed with 1x PBS pH 7.4 for three times and afterward incubated in a solution containing 4 % BSA (SEQENS) and 0.1 % Triton X-100 (Sigma Aldrich) for 1 hour to permeabilize the cell membrane and blocking. Before the staining procedure, samples were washed three times with PBS. Phalloidin 488 stain was incubated for 3 hours (RT) at a dilution of 1:1000 in PBS (Phalloidin-iFluor 488 Reagent, abcam, (ab176753)). Afterwards, samples were incubated with DAPI stain (1:500 in PBS) for 10 min (RT) and washed 4 times with PBS. Pluripotency staining for OCT4 was performed with the Oct3/4 primary antibody (Santa Cruz, sc-5279) at a concentration of 1:200 overnight at 4°C. Afterward, samples were washed 2 times with PBS and incubated with the secondary antibody Alexa Fluor^TM^ 488, 594, or 633, for 1 hour at RT with subsequent DAPI staining (above) or direct washing for 3 times with PBS. The germlayer-specific primary antibody stains were used in the following concentrations: Brachyury (RandD systems, AF2085) 1:20; GATA6 (Cell Signaling Technology, #5851) 1:1000; PAX6 (RandD systems, AF8150) 1:100. Cardiac-specific staining for cardiac Troponin T was performed with the primary cTnT antibody (biotechne, MAB18741-100) in a concentration of 1:200. Secondary antibodies and DAPI stains were applied as described before. Samples were stored in 1x PBS at 4 °C prior to imaging. Some samples were additionally incubated in a tissue clearing solution for 24 hours and imaged in the respective solution (Ultrapure water: 40 % D-Sorbitol, 10 % Glycerol, 4M Urea, 0.2 % Triton X-100, 15 % DMSO)[70]. All confocal images were recorded using the Opera Phoenix Confocal microscope (revvity).

#### Immunofluorescence staining (Photoreceptor progenitors)

Scaffolds were fixed with 4% paraformaldehyde at 4°C for 30 minutes. Samples were permeabilized with 0.3% Triton X-100 for 20 minutes at room temperature. 1% BSA in PBS was used for blocking. Samples were then incubated overnight at 4°C with primary antibody in 1% BSA solution containing 0.1% Triton X-100. After washing 3 times, 10 minutes each, the secondary antibody diluted in PBS containing 0.1% Triton X-100 solution was added to the samples and incubated overnight at 4°C. Samples were washed 3 times, 10 minutes each, before imaging with a laser scanning microscope. Primary antibodies used were: PAX6 (GenTex-Biozol GTX113241), OTX2 (R&D systems BAF1979), VSX2 (Sigma Aldrich AB9016), CRX (R&D systems, AF7085), Recoverin (Millipore, AB5585-1). Secondary antibodies used were: AF594 anti rabbit IgG (H+L) (Invitrogen A11012), AF594 anti sheep IgG (H+L) (Abcam AB150180), AF555 anti goat IgG (H+L) (Invitrogen (life technologies), A21432).

#### Clearing protocol (used for high-throughput experiments)

An alternative clearing protocol was applied for the automatically produced samples, since the lack of air flow inside the automated pipetting system does not allow for the use of hazardous chemicals. This protocol was adapted from the user protocol UP11 by ibidi^®^. After immunostaining was performed as described above, samples will be incubated in an increasing dilution series of Isopropanol, performed on ice. Add 30% Isopropanol (diluted with destilled water) and incubate the samples for 15 minutes on ice. Repeat this step consecutively with Isopropanol dilutions of 50% and 70%. Afterwards, incubate the samples in pure isopropanol for 15 minutes and repeat this step once. The samples have to be heated to room temperature afterwards, before the addition of Ethyl cinnamate (Sigma-Aldrich, CAS: 103-36-6). Incubate the sample in pure Ethyl cinnamate (99%) for 30 minutes to 1 hour before imaging. Samples will be imaged in Ethyl cinnamate.

#### Cryocutting and staining of slices

The samples were fixed in 4% PFA as described above. After washing with 1x PBS, samples were embedded in liquid OCT solution (Cellpath, KMA-0100-00A) in cryomolds (Sakura, 10x10x5mm³, Ref 4565). Samples were incubated in OCT for 1h before shock freezing. Shock freezing was done by placing a beaker with n-pentane into liquid nitrogen; afterwards, the cryomolds were placed into the n-pentane until the sample was completely frozen. Frozen cryomolds were stored overnight at – 20 °C. Cryocutting of samples was performed with a cryotome, Leica CM1950. The samples were cut into 5 µm slices throughout the whole sample to determine the middle area for analysis. Slices were stored on glass slides at -20 °C. Immunostaining was performed on the slices as follows: First, slices were placed in – 20°C cold acetone for 10 minutes, then they were dried and placed into MilliQ water for 10 min afterward. Staining afterwards was performed in a humidified petri dish using the same time points and concentrations as described above. After immunostaining, samples were mounted using DPX Mountant (Sigma Aldrich). Imaging was performed using the ECHO Revolution (ECHO a BICO company) microscope.

#### Live/Dead Staining of scaffolds

Viability assessment was performed after 2 days of culture (n=3) using the LIVE/DEAD Cell Imaging Kit (Invitrogen). The scaffolds were stained according to the protocol provided by the supplier and imaged using the Opera Phoenix Confocal microscope with a 10×/N.A. 0.3 air objective (revvity).

#### Image analysis

The image analysis of fluorescence intensity and maximum projection analysis was performed using the Harmony software (revvity). Further image analysis was conducted with the ImageJ software. Area analysis of scaffolds was performed, generating a 16-bit image, converting this image into a binary image using the default threshold method, and subsequently measuring the white pixel area. Diameter analysis was performed by applying the arrow tool for freehand measurements. Confocal image fluorescence intensity was analyzed by measuring the mean grey values of 16-bit converted maximum projection Z-Stack images.

#### Epigenetic analysis of germ layer differentiation

Pluripotent state and three lineage differentiation potential of iPSCs were analyzed by cell-type specific DNA methylation changes using PluripotencyScreen, as described in detail before [1]. The genomic DNA was isolated from the samples using the NucleoSpin Tissue Kit (Macherey-Nagel, 740952.50), and the yielded DNA was afterwards quantified with the NanoDrop 2000 spectrophotometer (Thermo Fisher Scientific). Bisulfite conversion of the genomic DNA was performed overnight using 500 ng DNA with the EZ DNA Methylation Kit (Zymo Research, D5002), and the elution was performed in 20 µL elution buffer. The PyroMark PCR Kit (Qiagen, ID. 978801) was used to amplify the target sequences using 2.5 mM Mg^2+^ and a primer concentration of 0.3 µm. Afterwards, targeted DNA methylation analysis was performed by pyrosequencing on a Q96PyroMark Q48 Autoprep Instrument (Qiagen) for three CG dinucleotides (CpGs) with characteristic modifications in the pluripotent state, mesoderm, endoderm, and ectoderm. The results can be used for deconvolution to estimate proportions of cellular differentiation toward these lineages. The primers for PCR and pyrosequencing can be found in Supplementary Table 2.

#### Metabolic activity assay

The metabolic activity analysis was performed using the alamarBlue™ reagent (Invitrogen^TM^) in the recommended concentration of 1:10 in iPS-Brew media. The samples (n=6) were incubated in 500 µL of the Resazurin solution for 2 hours at 37 °C in the incubator. Afterwards, 50 µL of the solution was transferred into a 96-well plate (n=3) and fluorescence was measured with a fluorescence plate reader (Tecan^TM^) according to the supplied protocol (emission: 590 nm; excitation: 530 nm).

#### RT-qPCR

Immediately after culture samples were lysed using the RLT-lysis buffer supplied with the RNeasy Mini Kit (Qiagen, 74104) and stored at -80 °C until further use. Prior to RNA-Isolation lysed samples were homogenized using QIAshredder columns (Qiagen, 79656) to remove the microgels from the lysis solution. Subsequently, RNA isolation was performed with the RNeasy Mini Kit (Qiagen, 74104), and concentration and purity were measured with NanoDrop™ UV/VIS Spectrophotometer. cDNA conversion was performed with 300-1000 ng of mRNA through application of the PrimeScript RT Master Mix (Takara Bio, RR036A) according to the protocol. RT-qPCR was performed with the SsoAdvanced Universal SYBR® Green Supermix (BioRad) on the CFX96 Touch Real-Time PCR Detection System (BioRad). As a reference gene, GAPDH was applied, and analysis was done using the 2^(-ΔCt)^- method. All utilized primers can be found in the Supplementary Table 1.

### RT qPCR (photoreceptor progenitors)

Total RNA was extracted using RNeasy kit (Qiagen). For each sample, 1 μg of RNA was reverse transcribed using a cDNA synthesis kit (Qiagen). All results were confirmed by three independent biological samples at each data point. Primers are listed in the supporting information Table 1.

#### Calcium imaging and video analysis of cardiac organoids

Calcium imaging was performed according to the protocol described by Y. Yao et al. [71]. Therefore, samples were stained with 4 µM Fluo4-AM diluted in 1x PowerLoad™ (Invitrogen^TM^, P10020) for 30 minutes. Afterward, samples were washed and calcium imaging was performed in calcium imaging buffer (in MilliQ water: 137 mM NaCl, 1.5 mM CaCl_2_, 5.4 mM KCl, 0.44 mM KH_2_PO_4_, 0.5 mM MgCl_2_, 0.4 mM MgSO_4_, 0.3 mM

Na_2_HPO_4_, 0.4 mM NaHCO_3_, 5.6 mM D-Glucose, 10 mM HEPES at pH 7.4). Calcium imaging analysis was performed using the NIS elements imaging software and the Nikon Eclipse Ti2-e microscope. The same microscope setup was used for the recording of the cardiac organoid beating videos.

#### Simulation of oxygen diffusion in scaffolds

Ansys CFX 2023 R1 was used to simulate oxygen transport within the organoids. This software uses the element-based finite volume method to solve numerically the governing equations. We idealized the organoid geometry as a fully spherical construct consisting of a continuum of cellular material and the microgels distributed within it. Stationary domains with no fluid flow were considered for both microgels and cells. The microgels’ positions within the organoids were generated by an IronPython script using normalized random number generation. The script checks if the distance between the microgels is greater than 2×r, where r is the radius of the microgel. For each condition, three randomly generated microgel distributions were obtained. The cells were modeled as a continuum material with defined cell density (1 × 10^14^ cells m^-3^), oxygen diffusion coefficient (1 × 10^-9^ m^2^ s^-1^), and oxygen consumption rate (2 × 10^-18^ mol cell^-1^ s^-1^ for iPSCs and 1 × 10^-17^ mol cell^-1^ s^-1^ for cardiomyocytes). The oxygen diffusion coefficient within the microgels was set to 2.5 × 10^-9^ m^2^ s^-1^. The oxygen concentration around the organoid was set to 0.22 mM. An isothermal condition was imposed, and the maximum root mean square (RMS) residual error of 10^−6^ was used as the convergence criteria for the transport equation. The grid study was performed to ensure that the proper element size was used, and a steady-state simulation was considered. Ansys CFX uses the equation below as the mathematical model for the simulation of oxygen transport within the stationary domains.

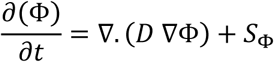

where Φ, 𝐷 and 𝑆_Φ_ are the oxygen concentration, kinematic diffusivity and volumetric source term.

### Statistical analysis

Statistical analysis and graphs were produced using Prism software. Box plots represent boxes and whiskers from minimal to maximal values, including all data points. The performed statistical analysis for each data set is reported in the respective figure caption. For statistical analysis, the P-values were represented in the figures.

## Supplementary figures

**Supplementary Fig. 1:**
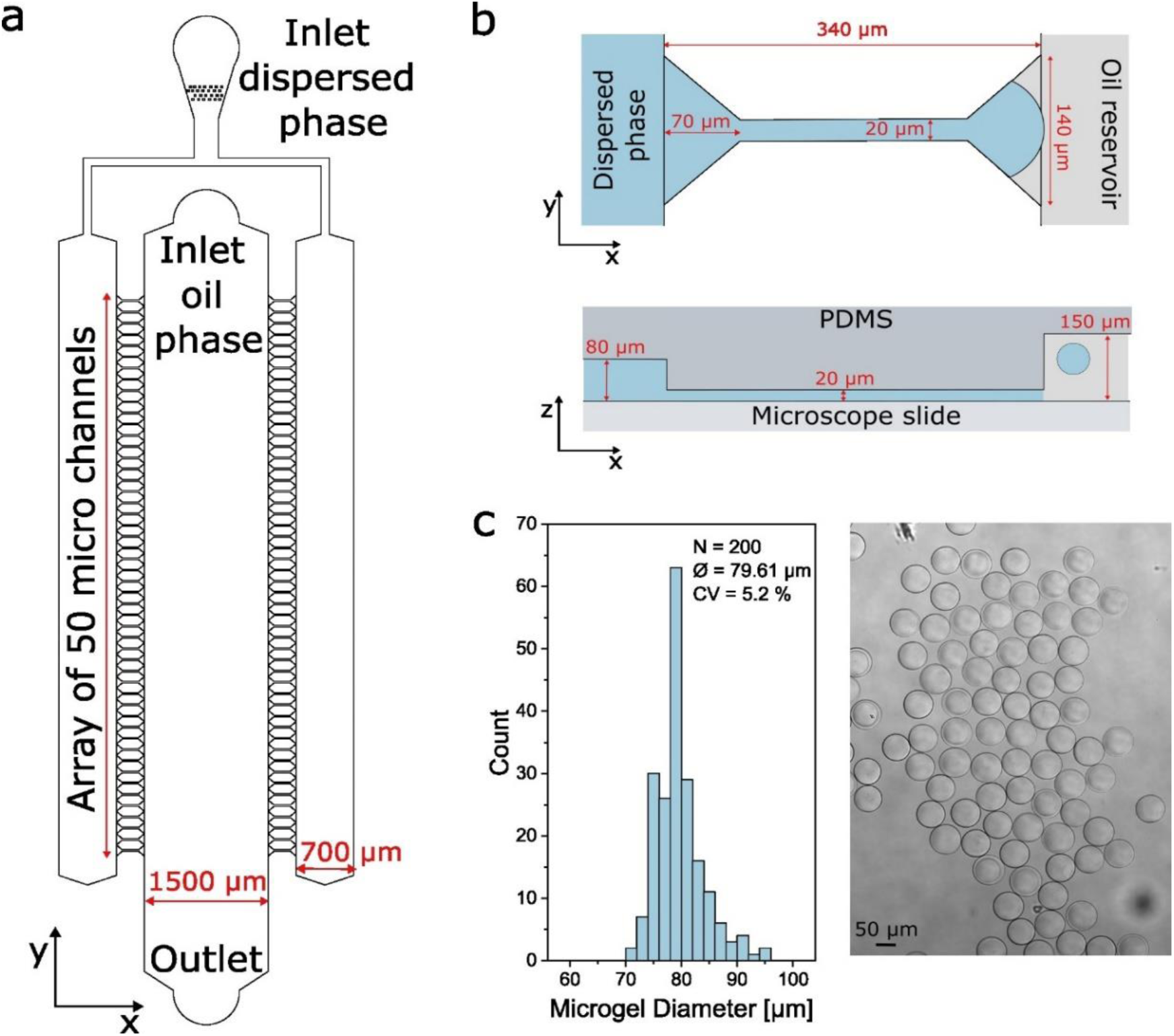
Characteristics of the microfluidic device: **a**, 2D design of the microfluidic step-emulsification device displaying the designated inlets and outlet as well as characteristic dimensions. **b**, Schematic of an individual micro channel with annotated dimensions. Top: top view on the micro channel and the reservoirs for the oil and dispersed phase. Bottom: side view on the micro channel highlighting the height differences between the different sections. **c**, Size distribution of PEG-DA-co-GMA microgels and microscope image in Milli-Q water after purification. The diameter of the microgels was measured by hand in Image-J. A total of 200 microgels were measured, resulting in an average diameter (Ø) of 79.61 µm, a standard deviation of 4.16 µm, and a coefficient of variation (CV) of 5.2%.

**Supplementary Fig. 2:**
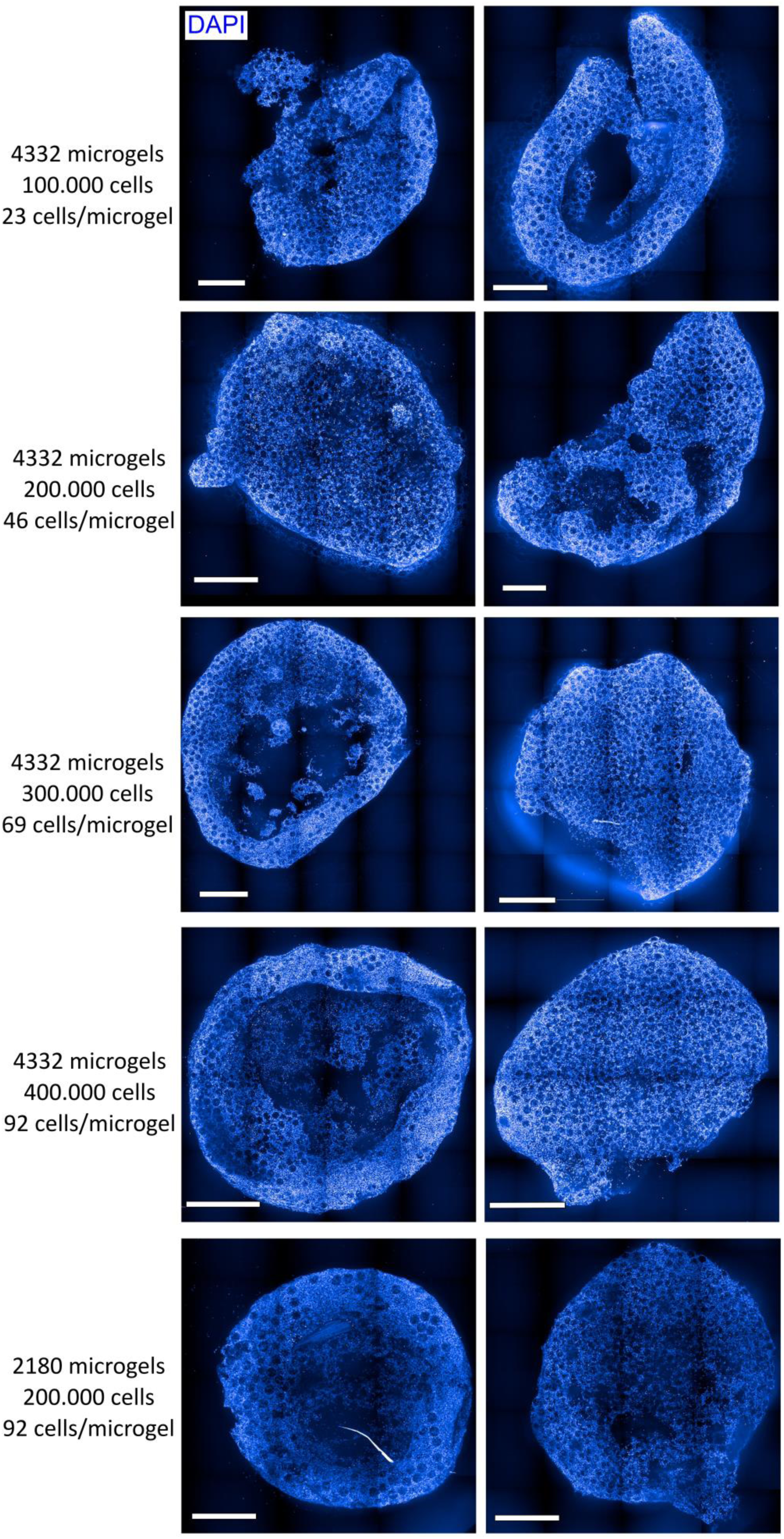
Optimization of cell/microgel ratio in mm-sized iPSC/microgel constructs to achieve shape and size consistency. Most promising conditions were 2180 microgels/200.000 cells and 4332 microgels/ 400.000 cells, which both represent the cell/microgel ratio of ∼92 cells/microgel. Blue: DAPI (nuclei); Scale bar: 1 mm.

**Supplementary Fig. 3:**
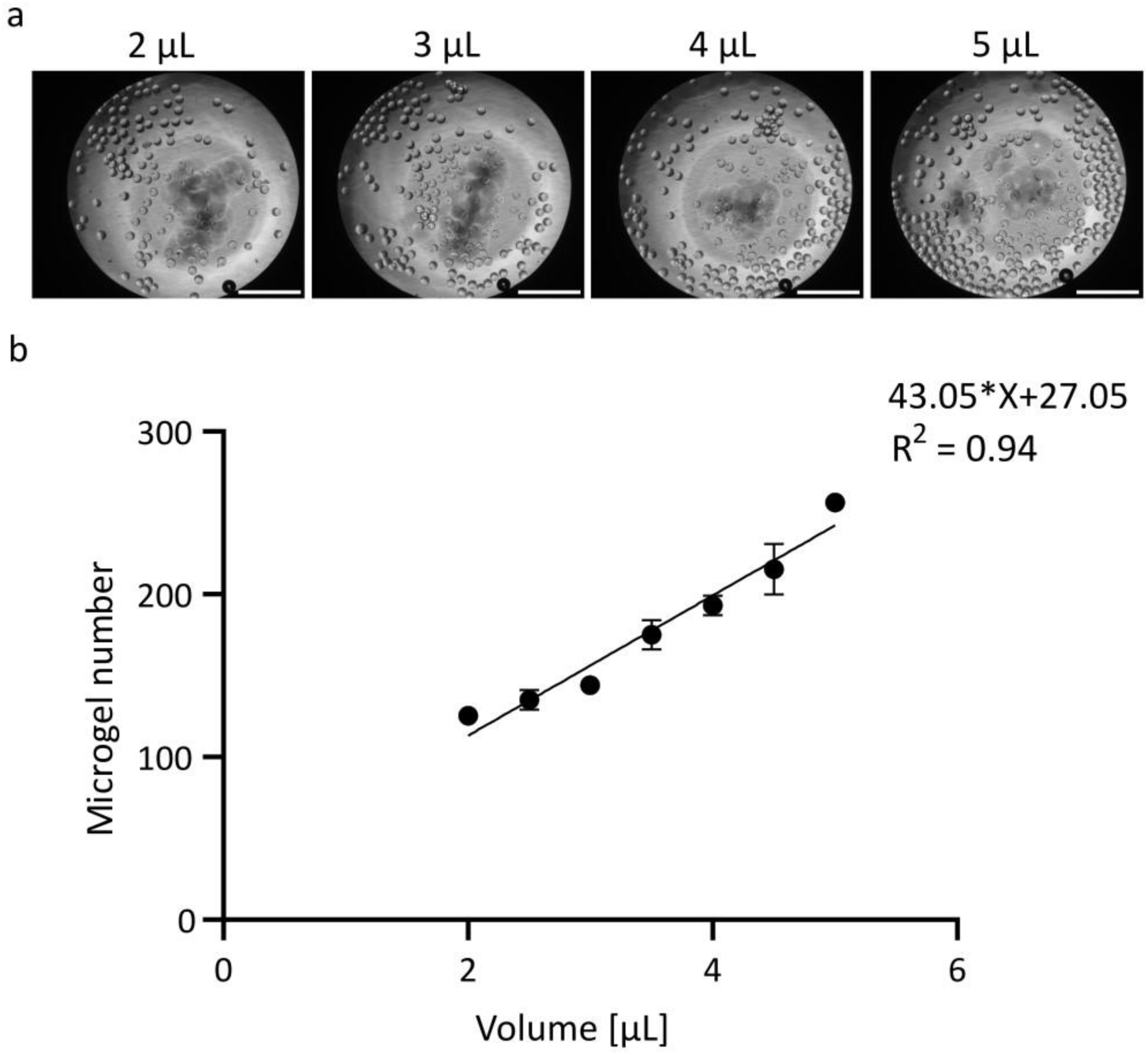
Calibration curve for the determination of microgel number in prepared microgel solutions. **a**, Brightfield images of different (2, 3, 4, 5 µL) volumes of microgel solution. Scale bar: 500 µm. **b**, Calibration curve of microgel number corresponding to microgel solution volume (each datapoint: n=3). Statistical analysis was performed with linear regression in Prism software.

**Supplementary Figure 4:**
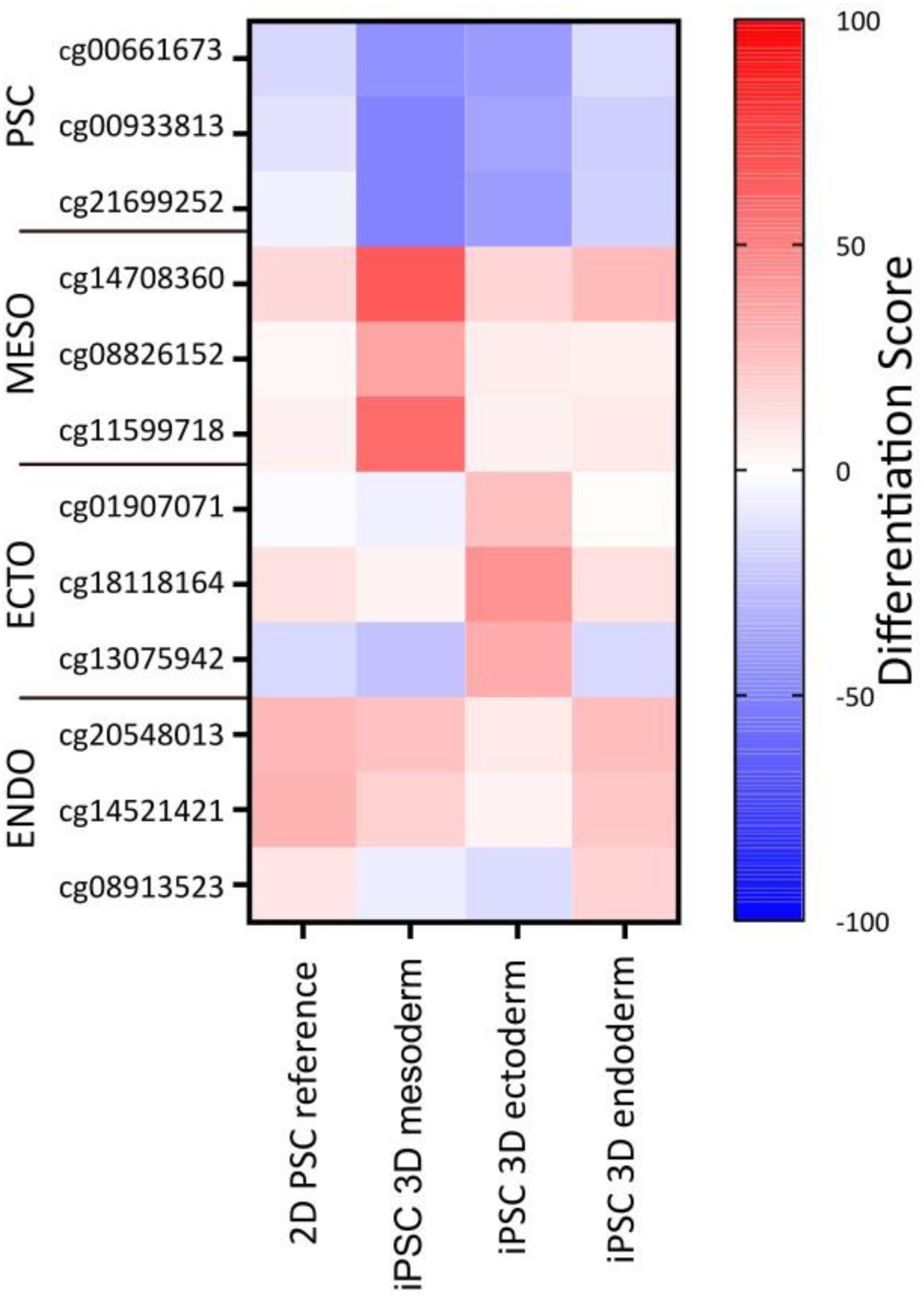
Methylation values for three distinct CpGs representing each germ layer. Methylation values were normalized to the iPSC reference for the iPSC cell lines UKAi009-A, UKAi0010-A, and UKAi0011-A (data from[1]). The heatmap shows the differentiation scores, which demonstrate the differences in the DNAm levels to the reference cell lines 102, 104, and 106 (regarding hypomethylated CpGs, 1 – DNAm was used as the calculation); In the heatmap, red indicates a change towards a certain methylation, while blue indicates the opposite. White demonstrates no change in comparison to the iPSC reference data.

**Supplementary Fig. 5:**
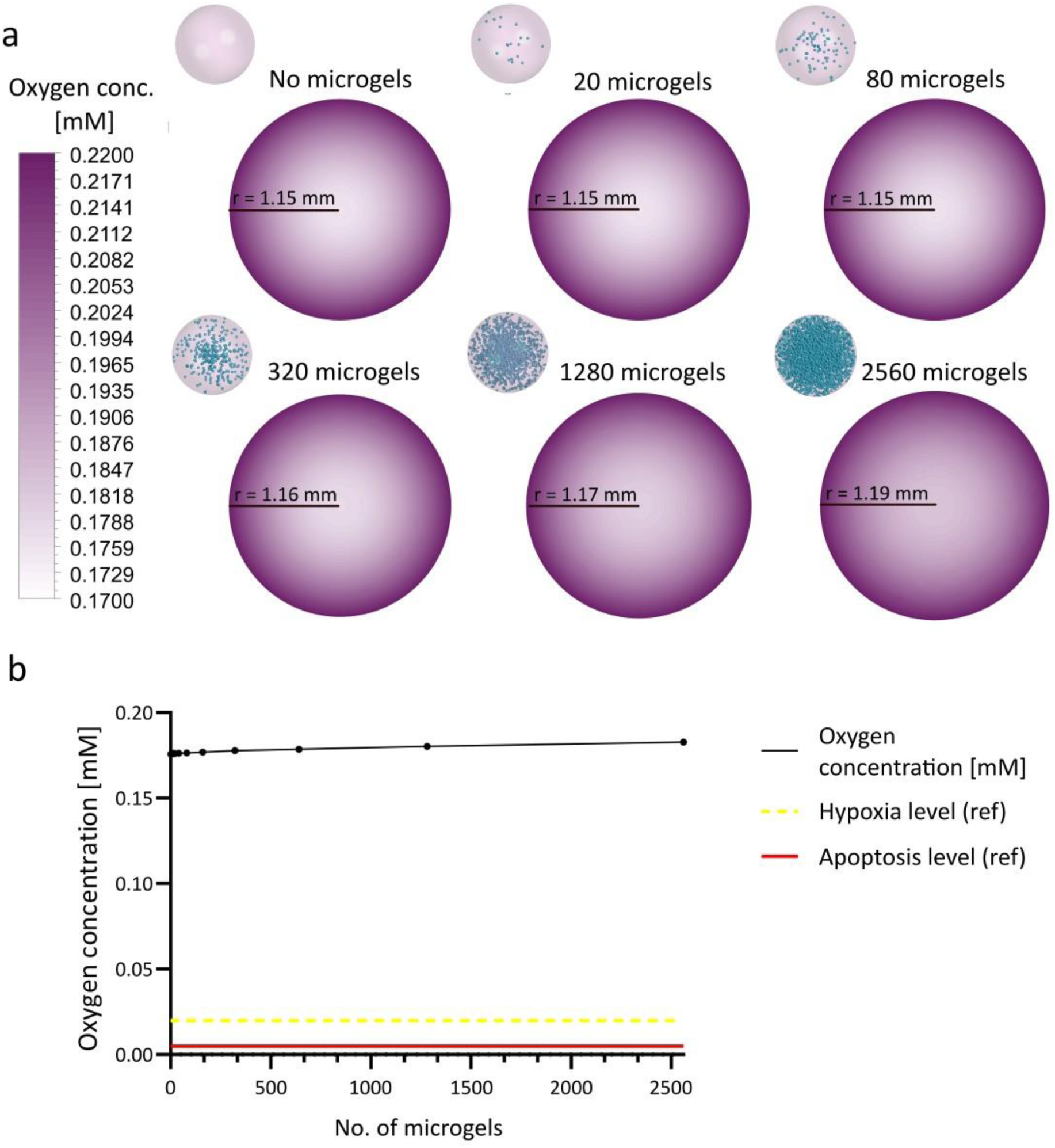
Simulation of oxygen diffusion in iPSC spheroid. **a**, Contour plots of oxygen concentration. Simulation of oxygen transport in a iPSC spheroid with a 1.15 mm radius does not show the formation of a necrotic core (C_Oxygen_ < 0.005 mM) and hypoxic area (0.005 mM < COxygen < 0.02 mM). **b**, Correlation between minimal oxygen concentration and number of microgels determined through simulation. Further details regarding oxygen transport modeling and simulation are reported in supplementary information.

**Supplementary Fig. 6:**
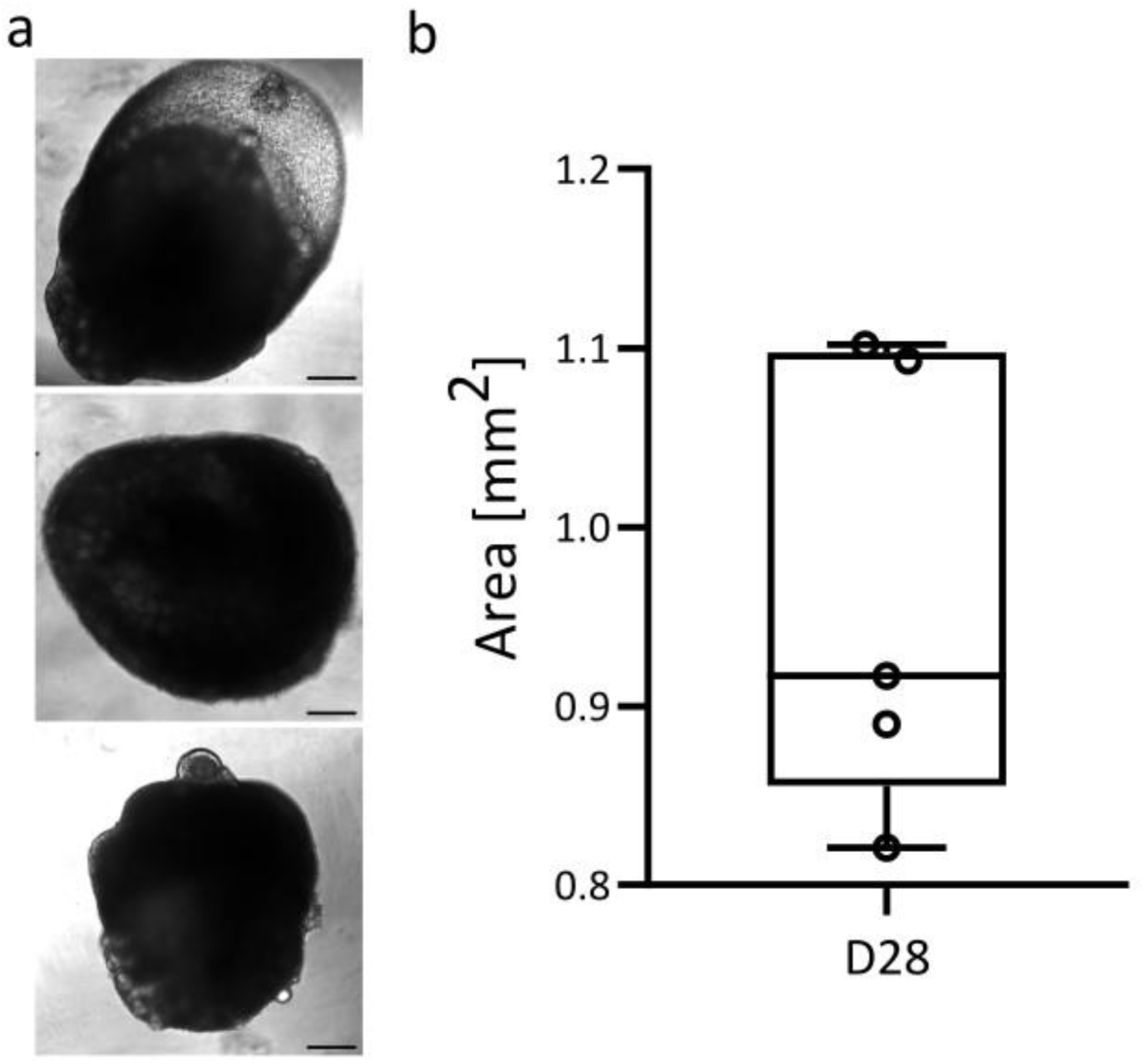
Area analysis of scaffolds produced with the 96-well format method. **a**, Brightfield images of three exemplary images of scaffolds on day 28 of culture used for the analysis in b. Scale bar: 200 µm. **b**, Area [mm^2^] and diameter [mm] were conducted with image analysis using ImageJ, n=6, statistical significance was calculated via unpaired t-test).

**Supplementary Fig. 7:**
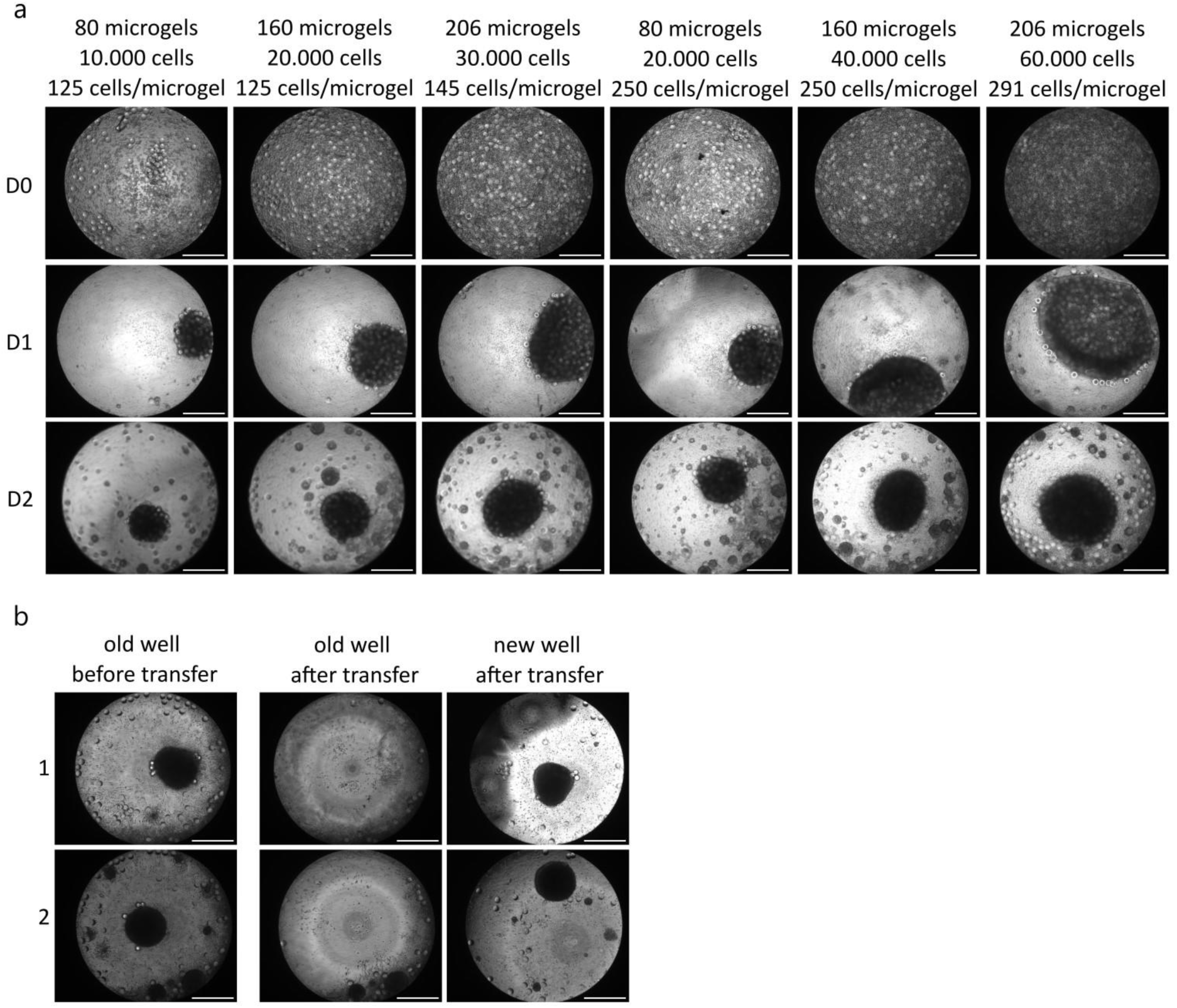
Pipetting analysis of microgels, cells and scaffolds with the automated liquid handling workstation JANUS G3. **a**, Scaffold generation in low-volume 384 well plates on day 0, 1 and 2 with different ratios and amounts of cells and microgels. Depending on the initial ratios and amounts used scaffolds of differing sizes were achieved. Scale bar: 500 µm. **b**, Transfer of scaffolds between wells using the automated pipetting on day 5. Successful transfer of scaffolds was observed resulting in removal of cell debris for further culture. Scale bar: 500 µm.

**Supplementary Fig. 8:**
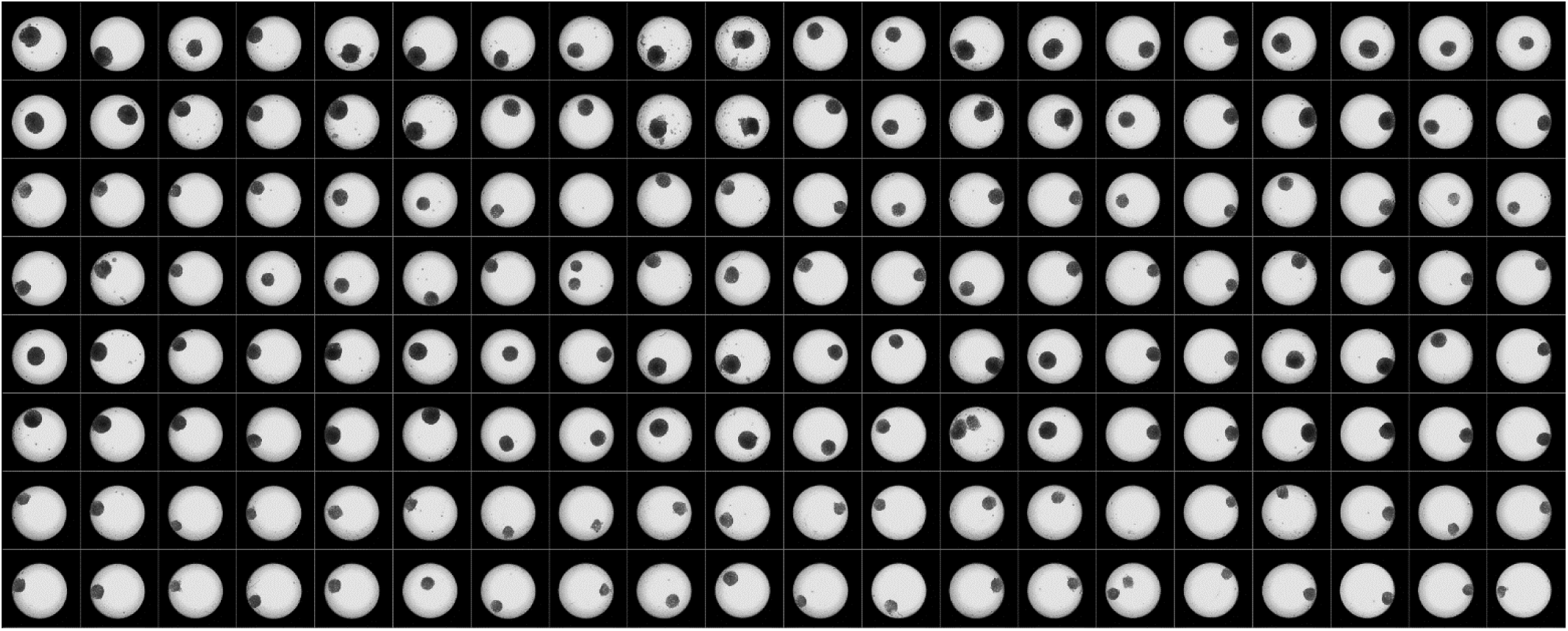
Representative view of 160 wells of a low-volume 384 well plate filled with iPSC/microgel scaffolds at day 2. Scaffolds were generated with the automated liquid handling system in a fully automated manner. This view includes different cell numbers between 5000 to 20.000 cells per well and different microgel amounts of either 80 or 160 microgels per well. Well diameter: 1.84 mm.

## Supplementary videos

**Supplementary Video 1**: Left: Microfluidic production of PEG-DA-co-GMA microgels. Right: Magnified video highlighting the droplet formation.

**Supplementary Video 2**: mm-scale cardiac organoid beating at D23 of differentiation. Uniform beating of the whole scaffold was observed. Video was recorded in real acquisition time.

**Supplementary Video 3**: Close-up of mm-scale cardiac organoid beating at D23 of differentiation. Tissue formation with embedded microgels was observed. Cardiomyocyte contraction was possible with microgels present in the tissue. Video was recorded in real acquisition time.

**Supplementary Video 4**: 96-well plate cardiac organoid beating at day 27 of differentiation. Uniform beating of the whole scaffold was observed. Video was recorded in real acquisition time.

**Supplementary Video 5**: Close-up of mm-scale cardiac organoid beating at D23 of differentiation. Tissue formation with embedded microgels was observed. Cardiomyocyte contraction was possible with microgels present in the tissue. Video was recorded in real acquisition time.

**Supplementary Video 6**: Formation of cell-assembled scaffolds in High-throughput production in low-volume 384-well plates. In this condition only cells were used and the assembly was demonstrated over 24 hours. Brightfield time lapse video. Diameter of well

**Supplementary Video 7**: Formation of cell-assembled scaffolds in High-throughput production in low-volume 384-well plates. In this condition 750 cells/microgel were used, and the assembly was demonstrated over 24 hours. Brightfield time-lapse video. Diameter of well 1.84 mm.

**Supplementary Video 8**: Formation of cell-assembled scaffolds in High-throughput production in low-volume 384-well plates. In this condition 375 cells/microgel were used, and the assembly was demonstrated over 24 hours. Brightfield time-lapse video. Diameter of well 1.84 mm.

**Supplementary Video 9**: Formation of cell-assembled scaffolds in High-throughput production in low-volume 384-well plates. In this condition 187.5 cells/microgel were used, and the assembly was demonstrated over 24 hours. Brightfield time-lapse video. Diameter of well 1.84 mm.

**Supplementary Video 10**: Formation of cell-assembled scaffolds in High-throughput production in low-volume 384-well plates. In this condition, 93.75 cells/microgel were used, and the assembly was demonstrated over 24 hours. Brightfield time-lapse video. Diameter of well 1.84 mm.

**Supplementary video 11**: Beating cardiac organoid at day 30. This organoid was generated from an iPSC/microgel scaffold with a ratio of 125 cells/microgel in a 384-well plate. The process was performed in an automated manner with automated liquid handling and a robotic arm. Diameter well: 1.84 mm.

## Supplementary table

**Supplementary Table 1:**
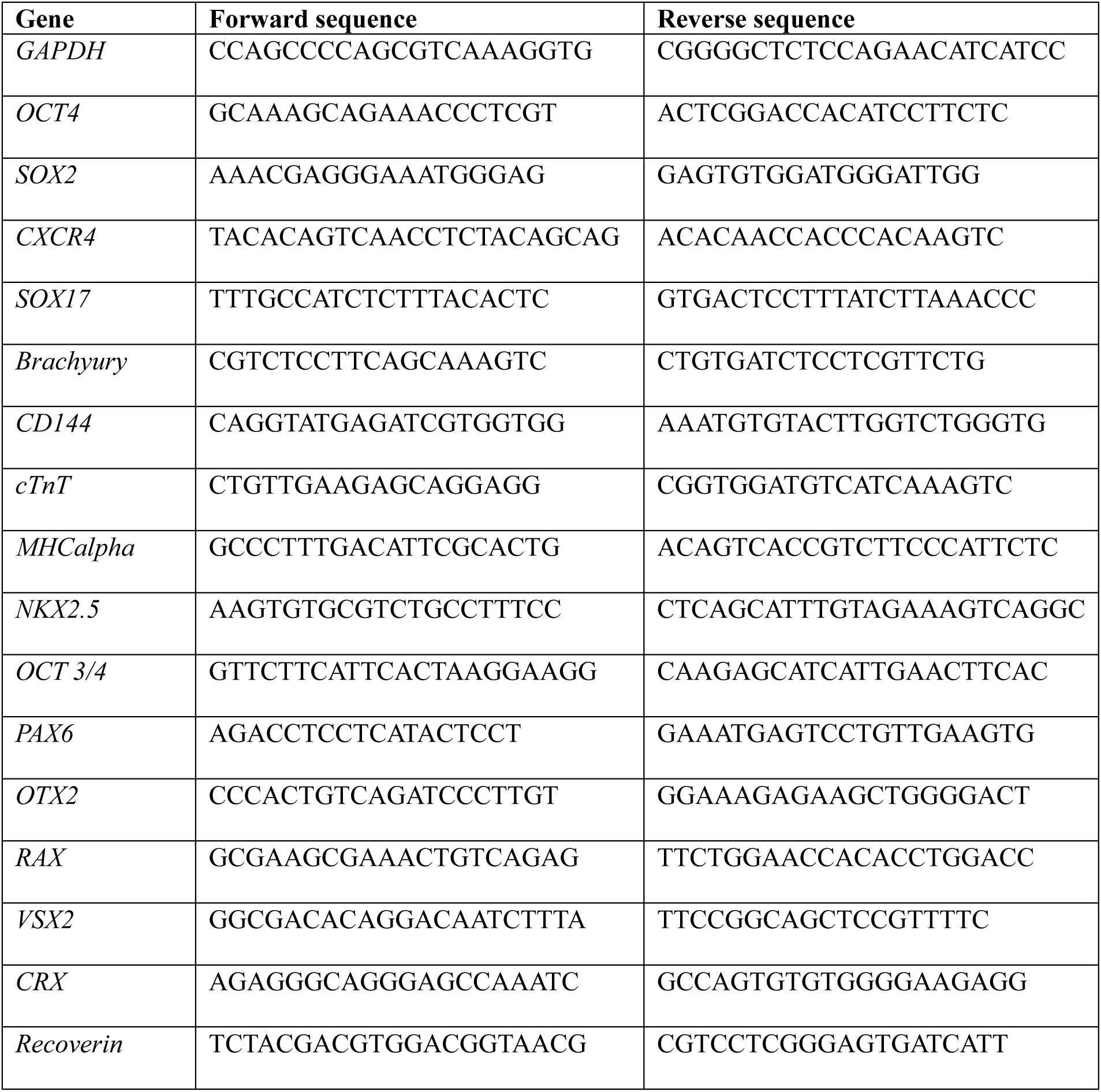
Primer used for RT-qPCR.

**Supplementary Table 2:**
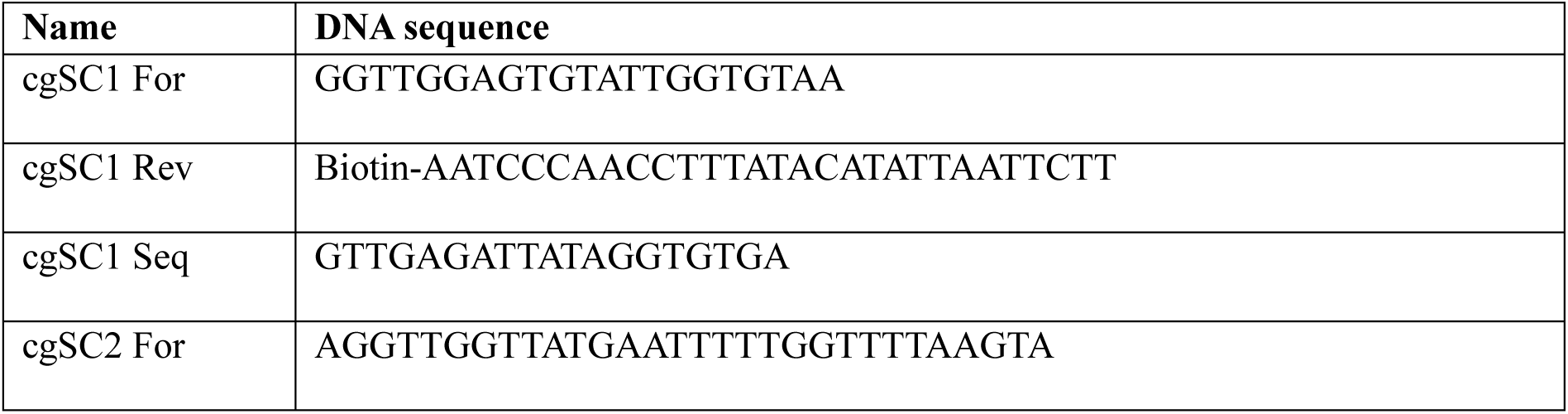
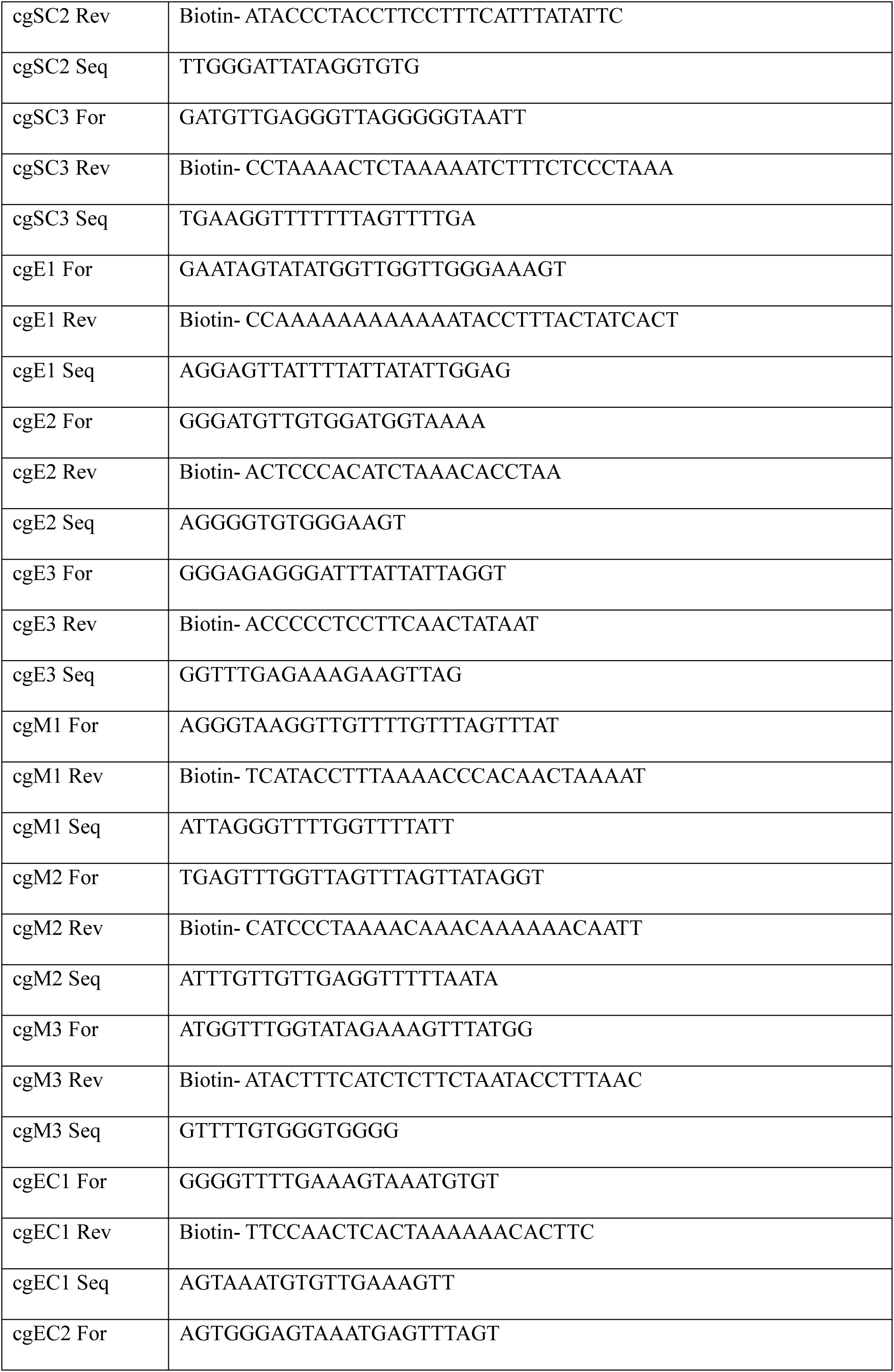
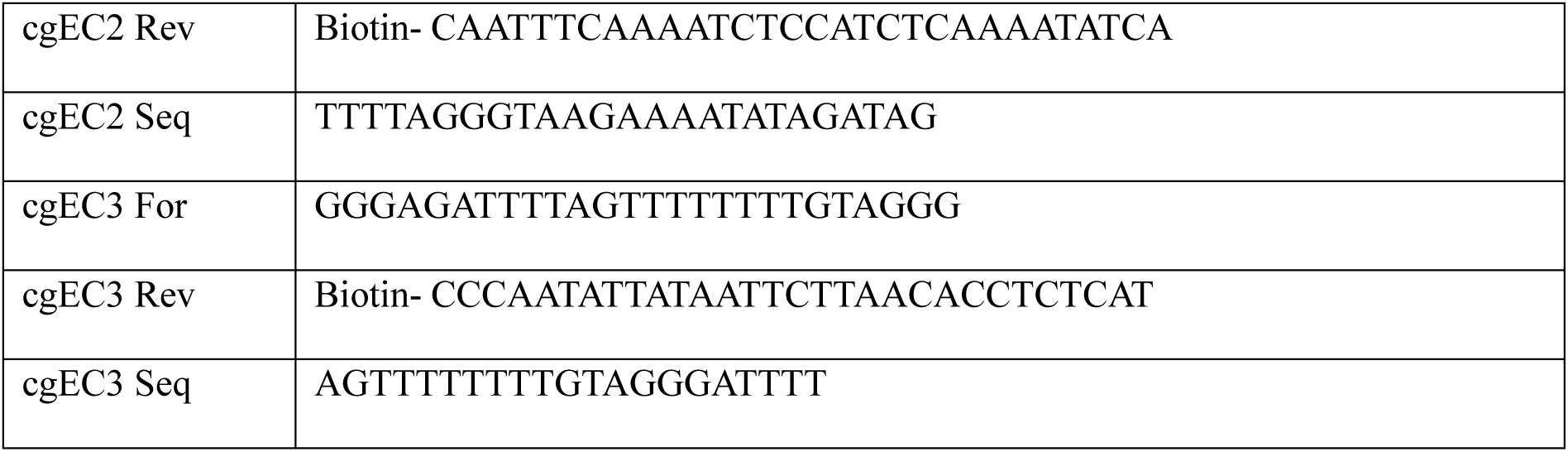
Primer for Pyrosequencing related to Figure 3.

## Acknowledgments

The authors gratefully acknowledge funding from the European Research Council within the ERC-2021-COG project 101043656, Heartbeat. We gratefully acknowledge funding from the German Research Foundation (DFG) within the project C3 and C9N of the Collaborative Research Centre (CRC 985) “Functional Microgels and Microgel Systems” and the Research Training Group Mechanobiology in Epithelial 3D Tissue Constructs (ME3T; 363055819/GRK2415). Furthermore, this work was supported through funding of the Senatsausschuss Wettbewerb (SAW) Leibniz-Transfer project µTISSUEfab and Leibniz Health Technologies Short Term Scientific Mission (STSM) funding, and by the Federal Ministry of Education and Research (VIP+: PluripotencyScreen; 03VP11580). The authors thank Greta Romahn for her contribution in preparing microgels via microfluidics, Paphassorn Karoon-ngampun for her help with the high-throughput experiment maintenance and preparation, and Céline Bastard for her help with cell culture and experiment maintenance.

